# Dietary derived Vitamin B12 protects the nematode *Caenorhabditis elegans* from Thiol Reducing agents

**DOI:** 10.1101/2022.05.03.490466

**Authors:** Alan D. Winter, Elissa Tjahjono, Leonardo J. Beltrán, Iain L. Johnstone, Neil Bulleid, Antony P. Page

**Affiliations:** Institute of Biodiversity, Animal Health and Comparative Medicine, University of Glasgow, Scotland, UK, G61 1QH; School of Life Sciences, University of Glasgow; Institute of Molecular, Cell and Systems Biology, University of Glasgow

**Keywords:** Vitamin B12, *Caenorhabditis elegans*, methyl transferase, Hypoxia, HIF-1, Redox, Thiol toxicity, methionine

## Abstract

We describe a novel SAM methyl transferase in the nematode *Caenorhabditis elegans* that is upregulated by thiol reducing agents and hydrogen sulfide with expression controlled by the hypoxia inducible factor pathway. This methyl transferase, RIPS-1, is expressed in the gut and hypodermis of this nematode with homologues found in a small subset of eukaryotes and bacteria, many of which can adapt to fluctuations in environmental oxygen levels. We identified RIPS-1 through forward genetic screens as the only gene that when mutated allowed worms to survive normally lethal concentrations of thiol reducing agents such as dithiothreitol (DTT) and β−mercaptoethanol. The RIPS-1 methyl transferase is an important player in the methionine cycle and its activation consumes methionine in a methionine synthetase and vitamin B12-dependant manner. This requirement limits the availability of vitamin B12 in the mitochondrion. Mitochondrial involvement was also established through a targeted enhancer screen that identified methylmalonyl-CoA epimerase as a strong genetic enhancer of RIPS-1 mutant resistance to DTT. Toxicity associated with thiol reducing agent exposure can be overcome in *C. elegans* by adding methionine, loss of RIPS-1, or by addition of excess vitamin B12. This work highlights the central importance of dietary vitamin B12 in normal metabolic processes in *C. elegans* and defines a new role in countering reductive stress

## INTRODUCTION

Cobalamin or vitamin B12 is made by a limited subset of prokaryotes but is required by a wide range of eukaryotes including the nematode worm *Caenorhabditis elegans* and mammals [1]. This essential vitamin exists in two biologically active forms, the mitochondrially required adenosyl cobalamin, a critical cofactor for methylmalonyl coenzyme A mutase (MMCM), and methyl cobalamin, an essential cytosolic cofactor for methionine synthetase [2]. These two functions are required in both humans and nematodes, particularly for the formation of methionine which is essential for providing methyl groups for numerous methylation processes.

The free-living nematode *C. elegans* acquires vitamin B12 from its bacterial food source with chronic B12 deficiency resulting in growth defects, infertility, and reduced lifespan [3]. In humans, vitamin B12 deficiency leads to megaloblastic anaemia and neurological problems [2]. In *C. elegans*, the cytosolic vitamin B12-dependant methionine synthetase enzyme METR-1 converts homocysteine to methionine, whereas the mitochondrial enzyme MMCM-1 converts methylmalonyl-CoA to succinyl-CoA, a reaction which follows an essential epimerase reaction catalysed by methylmalonyl-CoA epimerase (MCE-1).

The bacterial diet can have a profound effect on development of *C. elegans* and vitamin B12 has been shown to be an essential vitamin [4]. The standard *in vitro* food source of *C. elegans*, the *E. coli* strain OP50, is however mildly deficient in vitamin B12 when compared to HT115, the other commonly used *E. coli* food source, and other bacterial species such as *Comamonas* [4]. Mild vitamin B12 deficiency affects the conversion of methylmalonyl coenzyme A to succinyl-CoA in the mitochondrion, impairs the breakdown of propionate and branched chain amino acids, ultimately leading to mitochondrial dysfunction. A toxic build-up of propionate induces the gene *acdh-1* thereby activating a propionate shunt pathway, as shown in *C. elegans* using the transgenic *acdh-1*::*gfp* reporter which is a highly sensitive indicator of vitamin B12 deficiency [4].

Thiol reducing agents such as dithiothreitol (DTT) and β-mercaptoethanol represent powerful reducing agents that break disulfide bonds in proteins. While forward genetic screens have been performed to determine modes of resistance to oxidation [5], there is limited information regarding mechanisms of resistance to strong reducing agents. In this study we have applied the genetically amenable model nematode *C. elegans* to help elucidate mechanisms of resistance to strong reductive stress in a metazoan organism. This study revealed that resistance to thiol reducing agents is a vitamin B12 dependant process involving the methionine cycle in the cytosol and the TCA cycle in the mitochondria. Genetic screens for DTT resistance identified a single SAM methyl transferase, R08E5.3, loss of which affords protection from reductive stress. The Wormbase (https://wormbase.org) designation for R08E5.3 is RIPS-1 (for *Rhy-1*-Interacting Protein in Sulfide response) and we will use this gene name throughout. RIPS-1 expression is induced by thiol reducing agents and hydrogen sulfide (H_2_S). RIPS-1 induction is dependent on the transcription factor hypoxia inducible factor 1 (HIF-1) which represents the main oxygen sensing system in multicellular animals. Phylogenetic analysis reveals that RIPS-1 belongs to a group of methyl transferases found exclusively in metazoans including helminths and lancelets that can switch between aerobic and anaerobic metabolism.

## RESULTS

### Survival on the reducing agent DTT is dependent on bacterial food and growth media

Due to our long-standing interest in the control of cellular redox we examined the effect of thiol reducing agents such as dithiothreitol (DTT) on *C. elegans* development. During these experiments we made the unexpected observation that both bacterial diet and growth media had a dramatic influence on worm survival in the presence of DTT (Figure 1). On standard nematode growth media (NGM) wild type worms fed *E. coli* OP50-1, the most frequently used food source, stop development as larvae in the presence of 5 mM DTT (Figure 1B). In contrast, wild type worms fed with *E. coli* HT115(DE3) develop to fertile adults in the presence of the same concentration of DTT (Figure 1D), being almost indistinguishable from worms grown in the absence of DTT on either bacterial food source (Figure 1A and C). A major difference between the *E. coli* strains OP50-1 and HT115(DE3) is the level of vitamin B12 that these strains contain, and therefore provide as nutrient to *C. elegans*, with OP50-1 being relatively deficient in vitamin B12 compared to HT115(DE3) [6–8]. To examine this possible link between vitamin B12 and DTT resistance further we then tested the effect of different growth media. Standard nematode growth medium (NGM) uses peptone from animal sources and will contain vitamin B12, while NGM made from plant-based soybean peptone will be deficient in B12. The effect of media on DTT resistance was most noticible using *E. coli* HT115(DE3) (higher vitamin B12) where we found worm survival in the presence of DTT to be reduced with soybean peptone compared to standard NGM (Figure 1 D & H).

**Figure 1.**
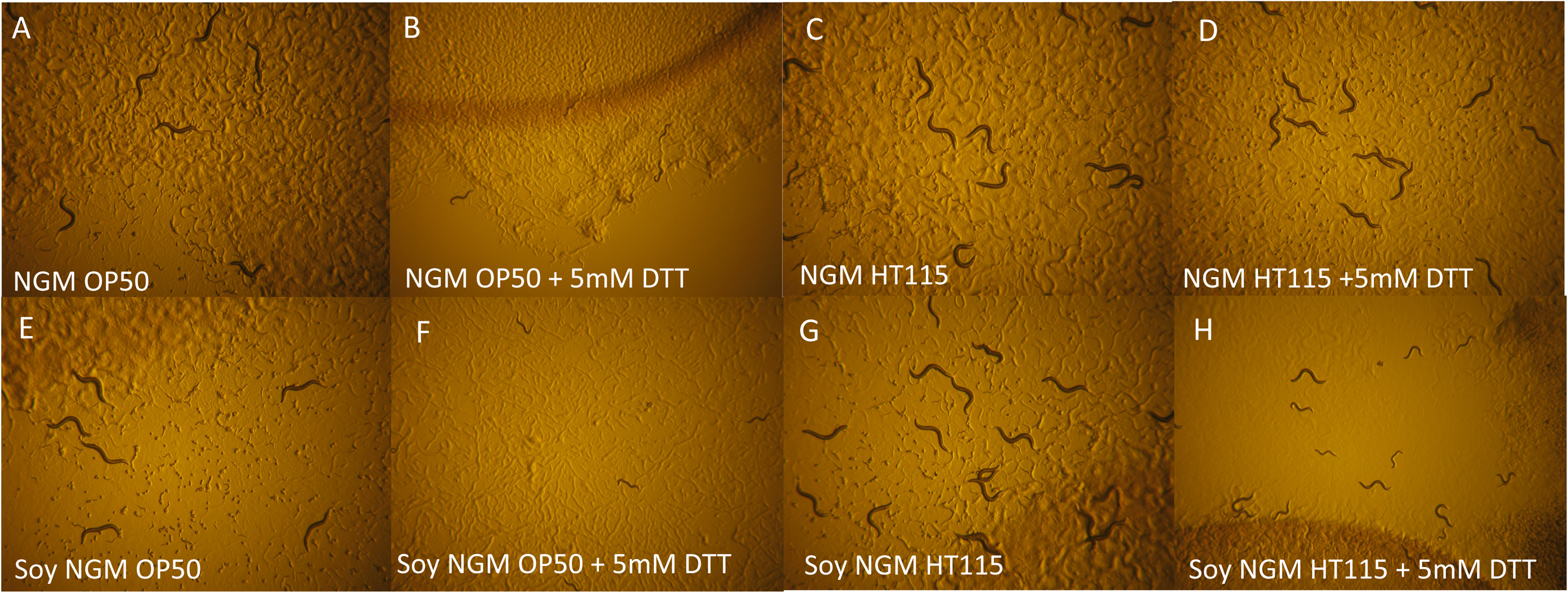
Survival on DTT is influenced by bacterial food and growth media. Plate images of *C. elegans* N2 on animal-derived peptone NGM agar and OP50-1 (A), with OP50-1 plus 5 mM DTT (B), with HT115(DE3) (C), with HT115(DE3) plus 5 mM DTT (D). On Soy-derived NGM agar with OP50 (E), with OP50-1 plus 5 mM DTT (F), with HT115(DE3) (G) and with HT115(DE3) plus 5 mM DTT (H). N2 Embryos added at day 0 and images take at day 4 at 10 x magnification.

### Supplementation with vitamin B12 results in METR-1-dependant survival on DTT

To define the role of vitamin B12 on survival of wild type worms under reductive stress we grew worms on soybean peptone NGM and *E. coli* OP50-1 (thereby providing *C. elegans* with low vitamin B12 from both growth media and food), and then compared worm growth on plates supplemented with cyanocob(III)alamin (vitamin B12) to those with no supplementation. Cyanocob(III)alamin is converted to both methylcobalamin, the cytoplasmic form of vitamin B12, and adenosylcobalamin, the mitochondrial form of vitamin B12. Figure 2C shows that supplementation with 64 nM vitamin B12 completely alleviated DTT sensitivity (Figure 2A). Vitamin B12 is used in only two enzymatic reactions; with cytoplasmic methionine synthetase (METR-1 in *C. elegans*) to convert homocysteine and methyl-tetrahydrofolate to methionine and tetrahydrofolate, and in the mitochondrial methylmalonyl-CoA mutase (MMCM-1) catalysed conversion of methylmalonyl-CoA to succinyl-CoA. To determine if the effect of vitamin B12 required either of these two enzymes we examined DTT survival using *metr-1* mutants and *mmcm-1* RNAi. *metr-1* mutants were susceptible to DTT toxicity even when supplemented with 64 nM vitamin B12 (Figure 2G), while with *mmcm-1* RNAi DTT toxicity was reversed on plates supplemented with 64 nM B12 (data not shown). Hence, DTT susceptibility is dependent on vitamin B12 and METR-1. Cobalamin compounds have been shown to be extremely effective in catalysing the autoxidation of sulfhydryl compounds to disulfides [9, 10]. This reported property of cobalamin could potentially provide a simple explanation for our results, with vitamin B12 essentially deactivating DTT. However, our finding that both vitamin B12 and active METR-1 are required to overcome the effects of DTT on development does not support this mechanism.

**Figure 2.**
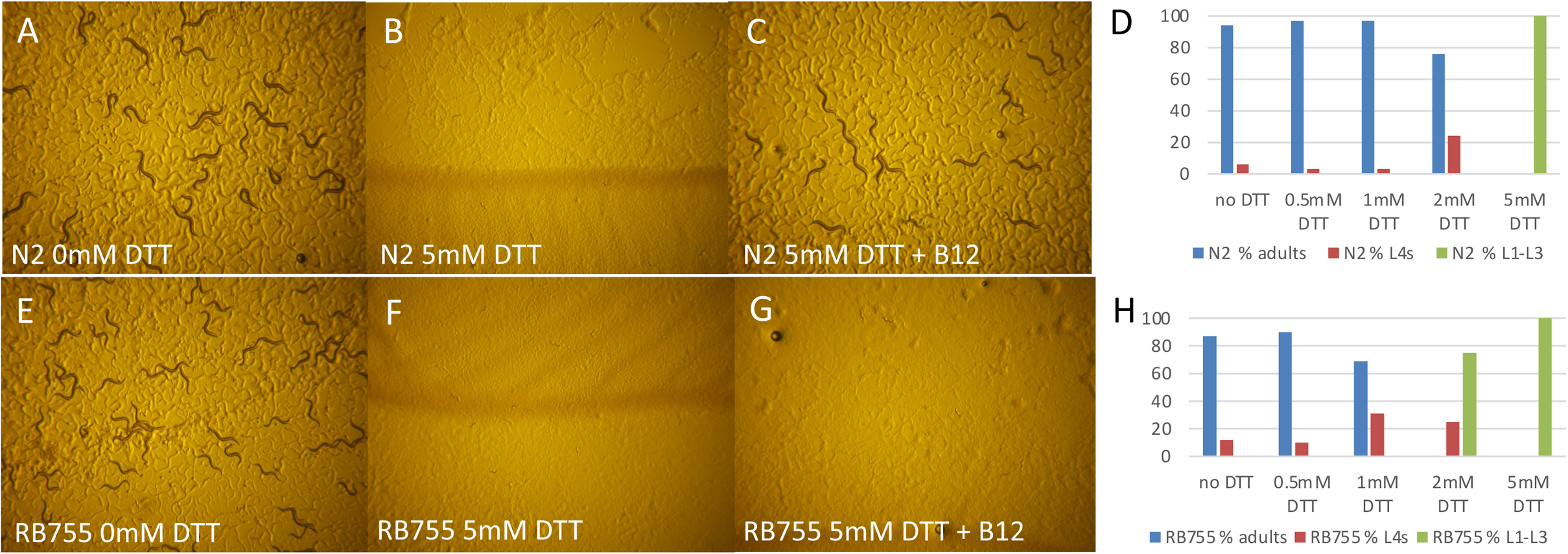
Supplementation with vitamin B12 results in METR-1-dependant survival on DTT. *metr-1* mutants are hypersensitive to DTT. Development of N2 *C. elegans* embryos over 4 days on soybean peptone NGM agar and *E. coli* strain OP50-1 in the absence of DTT (A), in the presence of 5 mM DTT (B) and in the presence of 5 mM DTT plus 64nM vitamin B12 (C). Plotted percentage development of wild type (N2) to adulthood after 4 days with no DTT (n=101), 0.5 mM DTT (n=119), 1 mM DTT (n=146), 2 mM DTT (n=145), and 5 mM DTT (n=114). Development of a *C. elegans metr-1* mutant [RB755, *metr-1*(*ok521*)] embryos over 4 days on soybean peptone NGM agar and *E. coli* strain OP50-1 in the absence of DTT (E), in the presence of 5 mM DTT (F) and in the presence of 5 mM DTT plus 64 nM vitamin B12 (G). Plotted percentage development of wild type (N2) to adulthood after 4 days with no DTT (n=71), 0.5 mM DTT (n=87), 1 mM DTT (n=67), 2 mM DTT (n=81) and 5 mM (n=58). All plate images at 5x magnification.

### Loss of METR-1 results in enhanced sensitivity to DTT

Due to the requirement of active METR-1 for vitamin B12-medited DTT resistance we looked to determine if a *metr-1* mutant displayed enhanced susceptibility to the effects of DTT. Titration of DTT from 0.5 to 5 mM demonstrated that the *metr-1* mutant strain is more sensitive to DTT (Figure 2H) compared to the wild type N2 strain (Figure 2D). At 2 mM DTT 76% (n=145) of wild type N2 but no *metr-1* mutants (n= 81) developed to adults.

### Loss-of-function mutations in the methyl transferase RIPS-1 confers resistance to DTT

A major advantage of the *C. elegans* model system is the ability to perform unbiased mutagenic screens. We reasoned that as DTT susceptibility is dependent on both vitamin B12 and on the B12-utilising enzyme METR-1, a screen aiming to identify DTT resistance mutants may uncover genes influencing either B12 availability, METR-1 function, or the methionine cycle. We therefore used EMS mutagenesis to isolate 14 independent strains that were resistant to 5 mM DTT and analysed nine of these using a combined SNP-based mapping and whole-genome re-sequencing protocol [11]. A strong genetic mapping signal to a 2-3 Mb region on the left-hand side of chromosome V was found for eight strains (Additional Data Figure 1A and B). Within this region the NGS data revealed that all eight strains contained homozygous variants for the methyl transferase *rips-1* (Figure 3 and Additional Data Table 2). We confirmed these eight alleles using conventional Sanger sequencing and then used Sanger sequencing only for the remaining five DTT-resistant alleles, all of which contained homozygous variants in *rips-1*. Therefore, from our mutagenesis screen we defined 13 *rips-1* alleles representing 11 unique variants, comprising six amino acid substitutions, four premature stop mutations, and a splice-acceptor site mutation (Figure 3 and Additional Data Table 1). We also analysed the *rips-1* allele *gk90219314* from the *C. elegans* Million Mutations Project [12]and confirmed it was also resistant to 5 mM DTT.

**Figure 3.**
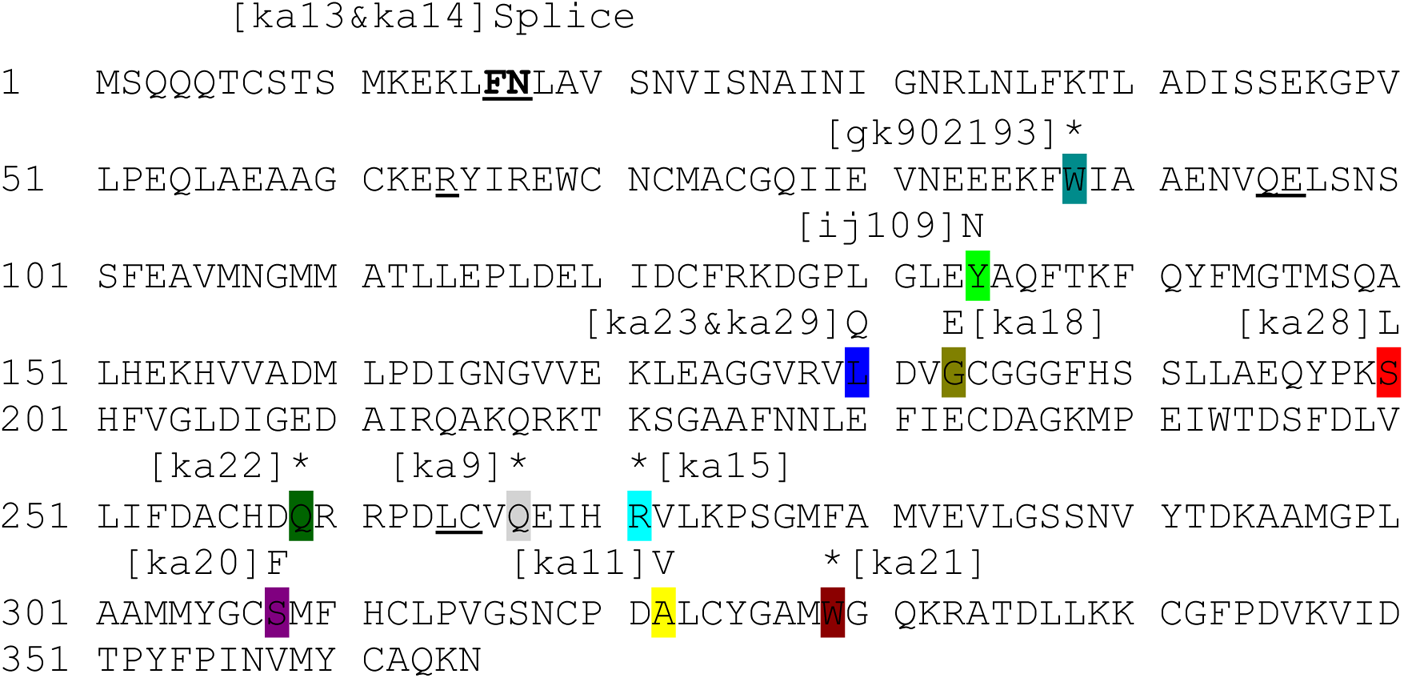
Protein sequence of RIPS-1 SAM methyl transferase highlighting location of mutation generated via EMS DTT resistance screen. Underlined residues exon/exon junctions and position and nature of mutation highlighted in colour and allele designation in brackets.

We then used RNAi with a 997bp *rips-1* fragment from the Ahringer lab *C. elegans* RNAi library [13] to confirm that *rips-1* was the gene conferring DTT resistance. We included three *rips-1* paralogues, *R08E5.1, R08F11.4,* and *K12D9.1*, identified by InParanoid 8 [14] and from our own BLAST analysis (Additional File 1). We performed two generation RNAi feeding, adding L4 larvae of the N2 wild type strain to RNAi bacteria and assessing the effects in the F1 progeny. Wild type worms treated with either *rips-1* RNAi or R08E5.1 RNAi survive well in the presence of 5 mM DTT whereas *R08F11.4* RNAi and *K12D9.1* RNAi did not (Additional Data Figure 2B). Because *rips-1* was identified as mutated in 13 independently obtained DTT resistant mutant strains, and in no case was R08E5.1 found to be mutated, it is most likely that the *R08E5.1* RNAi survival is the result of an off-target RNAi effect on *rips-1*. Accordingly, *rips-1* shares substantial nucleotide identity with the genomic fragment used for *R08E5.1* RNAi (89% overall identity, with multiple regions of 100% nucleotide identity (Additional Data Figure 2A). Finally, we introduced a wild type *rips-1* transgene into a *rips-1* mutant and found this restored DTT sensitivity (result described later and shown in Figure 9).

The resistance phenotype is depicted in Additional Data Figure 2B which shows that in the presence of 5 mM DTT no wild type N2 develop from embryos to adults in 4 days compared to a *rips-1* mutant, where almost all become adult. We quantified this survival at 0, 2.5, and 5 mM DTT. With no DTT, 88.5-95% of worms developed to adulthood for both strains. At 2.5 mM DTT, 17.7% of the wild type N2 strain develop to adults, while at 5 mM DTT no wild type worms reach adulthood, with all developmentally arrested as early larva (Figure 4A). For the *rips-1* mutants 97.8% and 88.7% of worms were adults after 4 days at 2.5 and 5 mM DTT respectively. We found similar levels of resistance for all our *rips-1*alleles (results not shown). Neither bacterial food nor growth media noticeably affect DTT resistance in a *rips-1* mutant (Additional Data Figure 3). We then tested if the resistance found in our mutants extended to other thiol reducing agents. In the presence of 2.5 mM β-mercaptoethanol 84% of *rips-1* mutants develop to adults compared with 2.6% of wild type N2 (Figure 4B).

**Figure 4.**
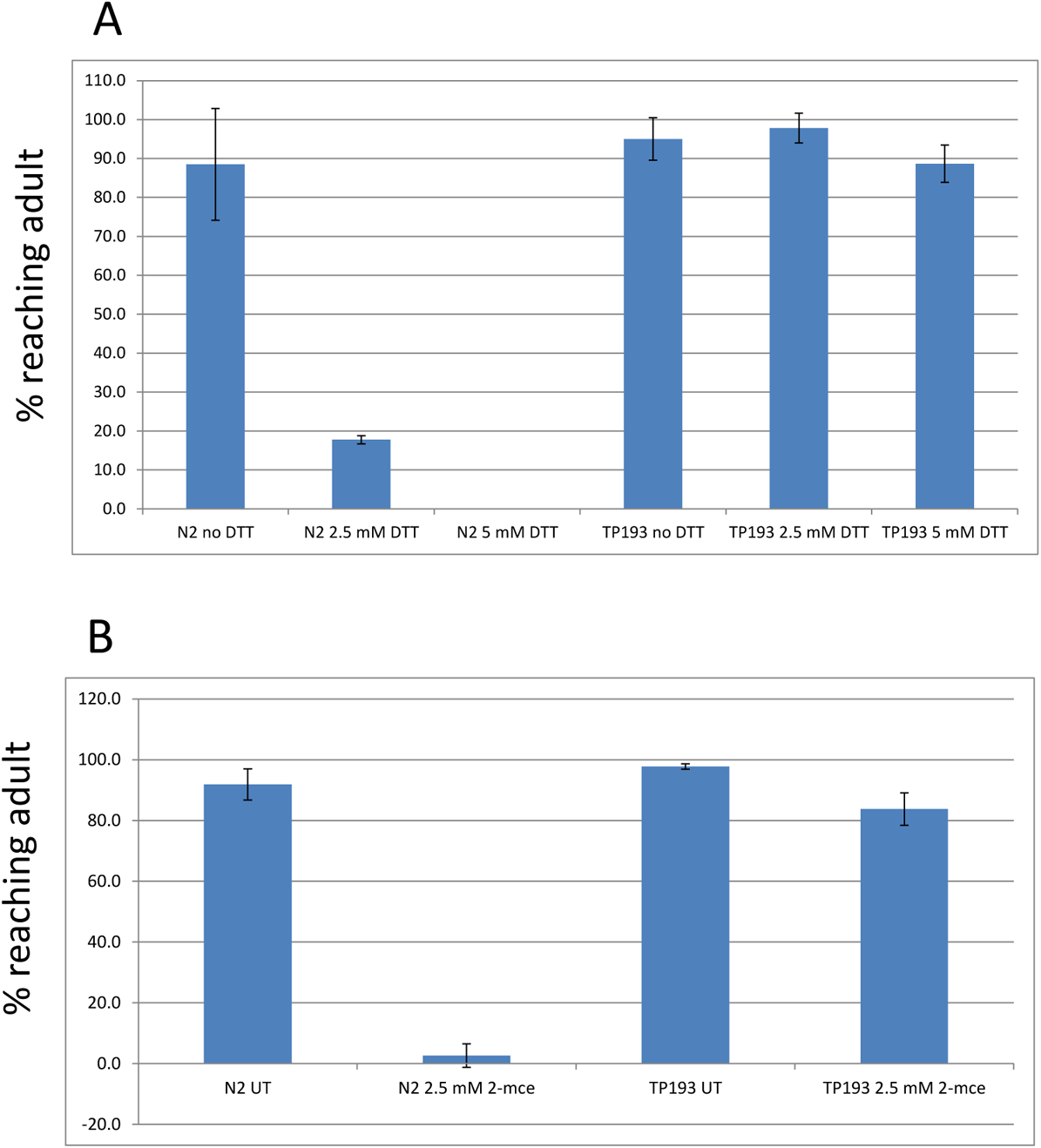
*rips-1* mutants are highly resistant to thiol reducing agents. Embryos of wild type (N2) and DTT resistant *C. elegans* [TP193, *rips-1*(*ij109*)] were place on Soy peptone NGM agar plates with OP50-1 and percentage survival to adulthood was determined after 4 days on plates supplemented with 0 mM DTT, 2.5 mM DTT and 5 mM DTT (A) and likewise on 0 mM 2-mercaptoethanol and 2.5 mM 2-mercaptoethanol (B).

### RIPS-1 is a methyl transferase conserved in diverse species but not present in vertebrates

To gain more insight into the role of RIPS-1, beyond its presumed biochemical activity as a methyl transferase, we identified proteins related to RIPS-1 and evaluated previous functional studies. Methyl transferases represent a large gene family, with 208 genes identified in humans [15]. BLAST analysis generated multiple hits based on the methyl transferase domain (pfam 13847: corresponding to residues 175-283 of RIPS-1). For our analysis we therefore required BLAST hits to have ≥30% identity over ≥90% of the full 365 amino acid residues of RIPS-1 (Additional Data File 1). Using the taxids described in Additional File 1, this analysis identified related proteins in multiple species of free-living and parasitic nematodes (round worms such as *C. elegans*), bacteria, mycobacteria, archaea, a single species of fungi (*Arthrobotrys oligospora*), and in multiple non-nematode multicellular organisms, such as *Branchiostoma belcheri* (lancelet), *Strongylocentrotus purpuratus* (crown-of-thorns starfish) and *Strongylocentrotus purpuratus* (purple sea urchin). However, these genes remain uncharacterised in these species. We noted that the RIPS-1 residues involved in the six amino acid substitution mutations are well conserved in these putative orthologs (Additional Data Figure 4). Likewise, these residues are also present in the paralogous *C. elegans* proteins (Additional Data Figure 4). Based on our criteria we found no proteins related to RIPS-1 in any plant, insect, platyhelminth (flat worms), annelida (segmented worms), or vertebrate species (Additional Data File 1). We also searched for remote homologues using HMMER hmmsearch[16] using the proteins aligned in Additional Data Figure 4A as input but found no likely RIPS-1 orthologs in vertebrates. In support of this, OrthoList [17], which identifies the likely human orthologs of *C. elegans* genes using a meta-analysis of four orthology-prediction methods, does not show an orthologous match in humans for *C. elegans* RIPS-1.

### DTT resistance due to loss of RIPS-1 is not dependent on METR-1

We had established two conditions that caused DTT resistance; high levels of vitamin B12 in combination with active METR-1 (methionine synthetase), and loss-of-function mutations in RIPS-1. To test if the mechanism by which loss of RIPS-1 caused DTT resistance was dependant on active METR-1, we constructed an *rips-1*; *metr-1* double mutant and compared survival of *rips-1* and *metr-1* single mutants with the double mutant on 5 mM DTT (Figure 5). The *rips-1*; *metr-1* double mutant showed comparable resistance to the *rips-1* single mutant, with 77% embryos developing to the adult in the presence of 5 mM DTT compared with 90% for the *rips-1* single mutant. This contrasts to no embryos developing to adults on 2 or 5 mM DTT for the *metr-1* single mutant (not shown). Therefore, the mutant *rips-1* DTT resistance phenotype is not suppressed by loss of *metr-1*, indicating that the mechanism by which loss of RIPS-1 confers resistance does not require active METR-1.

**Figure 5.**
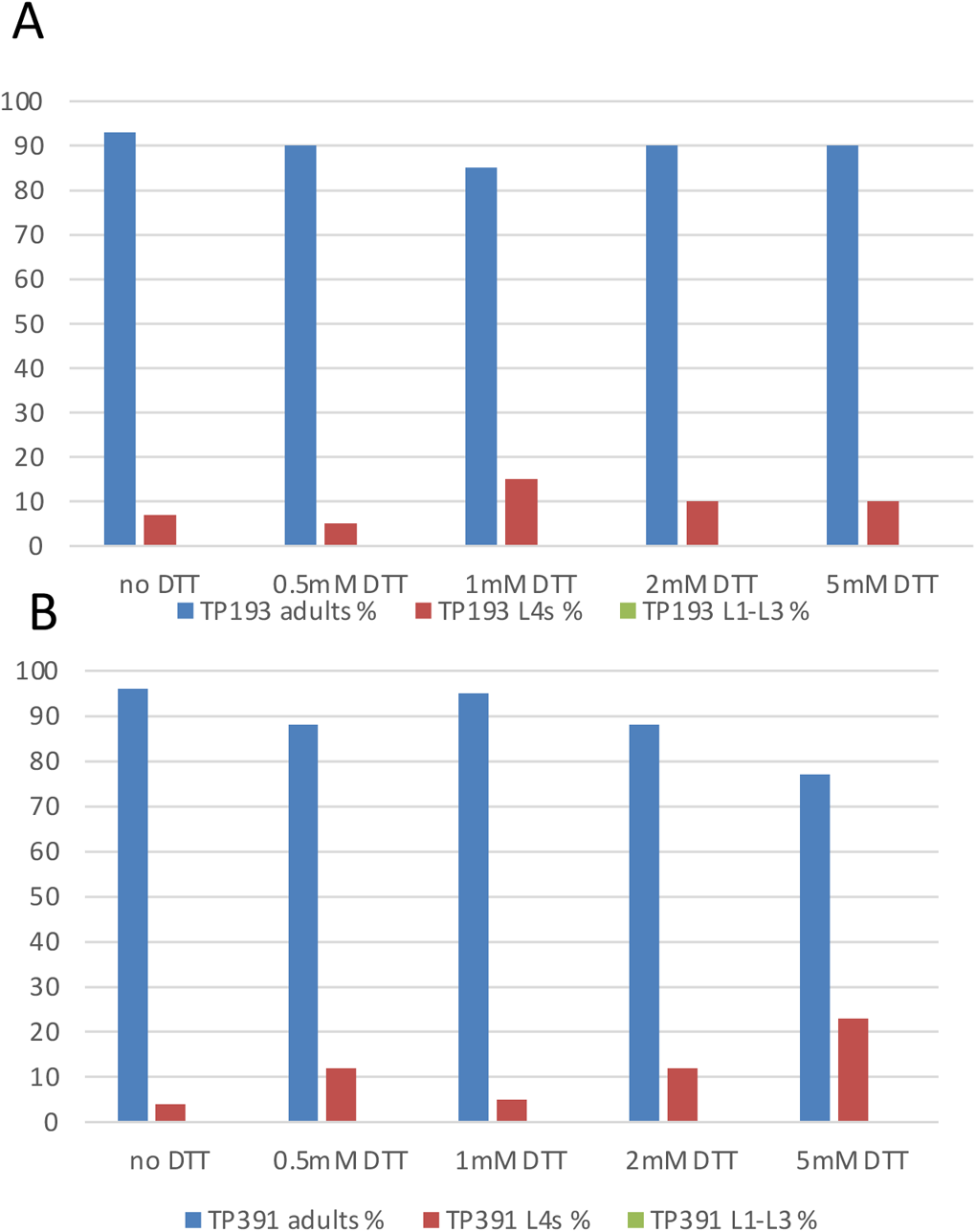
DTT resistance due to loss of *rips-1* is not dependent on METR-1. *rips-1* mutant embryos [TP193, *rips-1*(*ij109*)] (A) were assessed for development to adults after 4 days on 0 mM DTT (n=102), 0.5 mM DTT (n=100), 1 mM DTT (122), 2 mM DTT(88) and 5 mM DTT(n=87) and compared to (B) *metr-1;rips-1* double mutant [TP391 *metr-1*(*ok521*); *rips-1*(*ij109*)] on 0 mM DTT (n=140), 0.5 mM DTT (n=136), 1 mM DTT (n=119), 2 mM DTT(n=135) and 5 mM DTT(n=105).

### Combined loss of *rips-1* and *mce-1* results in enhanced DTT resistance

Due to the role established for vitamin B12 and METR-1 (methionine synthetase) we performed a targeted RNAi modifier screen aiming to identify additional genes that might suppress or enhance DTT sensitivity in a *rips-1* mutant. *C. elegans* genes involved in B12 processing, the methionine cycle, the folate cycle, the transulferation pathway, AdoCbl processing, the propanoic acid pathway and the propanoic acid shunt were identified from a review of the literature [18–23]. RNAi clones for the *C. elegans* genes targeted along with pathways and gene functions are listed in Additional Data Table 1. RNAi was performed in the *rips-1* mutant strain in the presence and absence of DTT. In the absence of DTT none of the 24 genes tested resulted in any additional phenotypes in the *rips-1* mutant background in comparison to the wild type N2 background (not shown). Testing of the 24 gene mini-RNAi library in the presence of DTT led to the identification of a strong enhancer of *rips-1* DTT resistance, the methylmalonyl-CoA epimerase gene (*mce-1*) (Figure 6B). None of the other 23 genes tested reproducibly altered the DTT resistance phenotype. We confirmed the *mce-1* enhancer result by constructing an *mce-1;rips-1* double mutant (Figure 6E & F). Importantly, the *mce-1* single mutant confers no resistance to 5 mM DTT, being as susceptible as wild type worms to this reducing agent (not shown). In combination however, *rips-1* and *mce-1* mutations confer high level of resistance to DTT and survive 7 mM DTT (Figure 6F), compared to a maximum survival concentration of 5 mM for the *rips-1* single mutant (Figure 6C).

**Figure 6.**
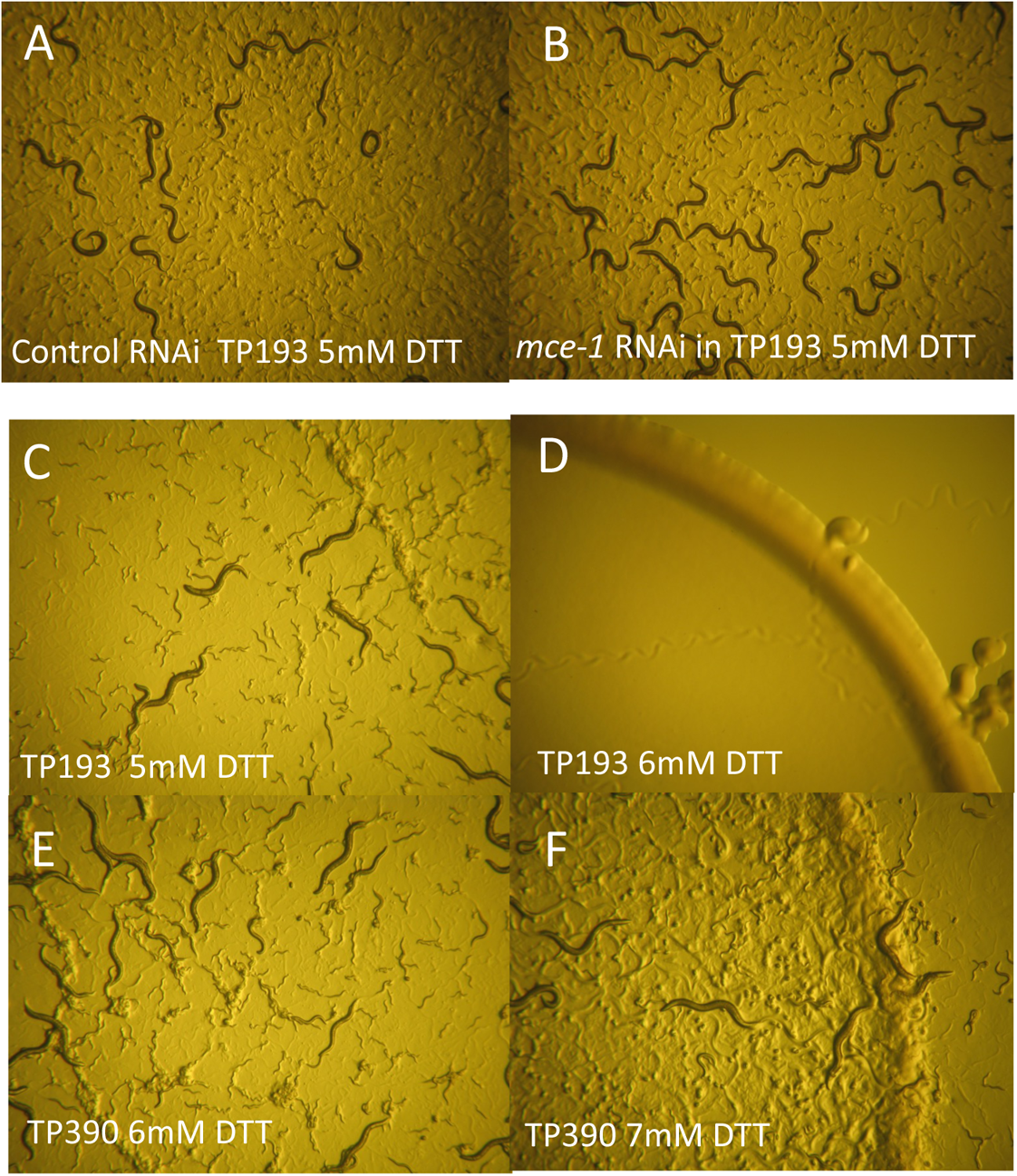
Genetic interaction screen identified *mce-1* (methylmalonyl-CoA epimerase) as an enhancer of DTT resistance in the *rips-1* mutant background. RNAi by feeding was performed in a *rips-1* mutant strain [TP193, *rips-1*(*ij109*)] with control (vector only RNAi L4440) (A) or D0203.5 (*mce-1*) RNAi (B) followed by selection on 5 mM DTT plates. Enhanced resistance was confirmed for a double mutant on DTT, as the *rips-1* single mutant [TP193, *rips-1*(*ij109*)] survived 5 mM DTT (C) but was killed on 6 mM DTT (D). The *mce-1;rips-1* double mutant [TP390, *mce-1*(*ok243*); *rips-1*(*ij109)*] survived well on both 6 mM DTT (E) and 7 mM DTT (F). 10x magnification.

### Methionine supplementation partially suppresses susceptibility to DTT

METR-1 (methionine synthetase) utilises vitamin B12 in the conversion of homocysteine and methyl-tetrahydrofolate to methionine and tetrahydrofolate. We wondered therefore if one, or both, of the products of this reaction might influence the DTT resistance phenotype. We found tetrahydrofolate highly insoluble above 100 µM in our plate supplementation assays, and therefore restricted our analysis to methionine. Development of L1 larvae placed on 5 mM DTT plates in the presence of methionine at 0, 1, 10 and 20 mM was examined (Figure 7). Without methionine supplementation no development to adulthood was observed for wild type (N2) (n= 166) or the *metr-1* single mutant (n= 217), while at 20 mM methionine 37% (n=119) and 30% (n=143) were adults after 4 days respectively. For the *rips-1* single mutant addition of 20 mM methionine did not alter the resistance phenotype, while in the *rips-1*; *metr-1* double mutant a reduction from 99% adults at 0 mM (n= 303) to 75% adults at 20 mM (n=89) was observed (Figure 7). This result shows that in the presence of RIPS-1, methionine supplementation results in partial DTT resistance, but when RIPS-1 is absent supplementation has no effect. This suggests that both may influence resistance to DTT via the same molecular mechanism.

**Figure 7.**
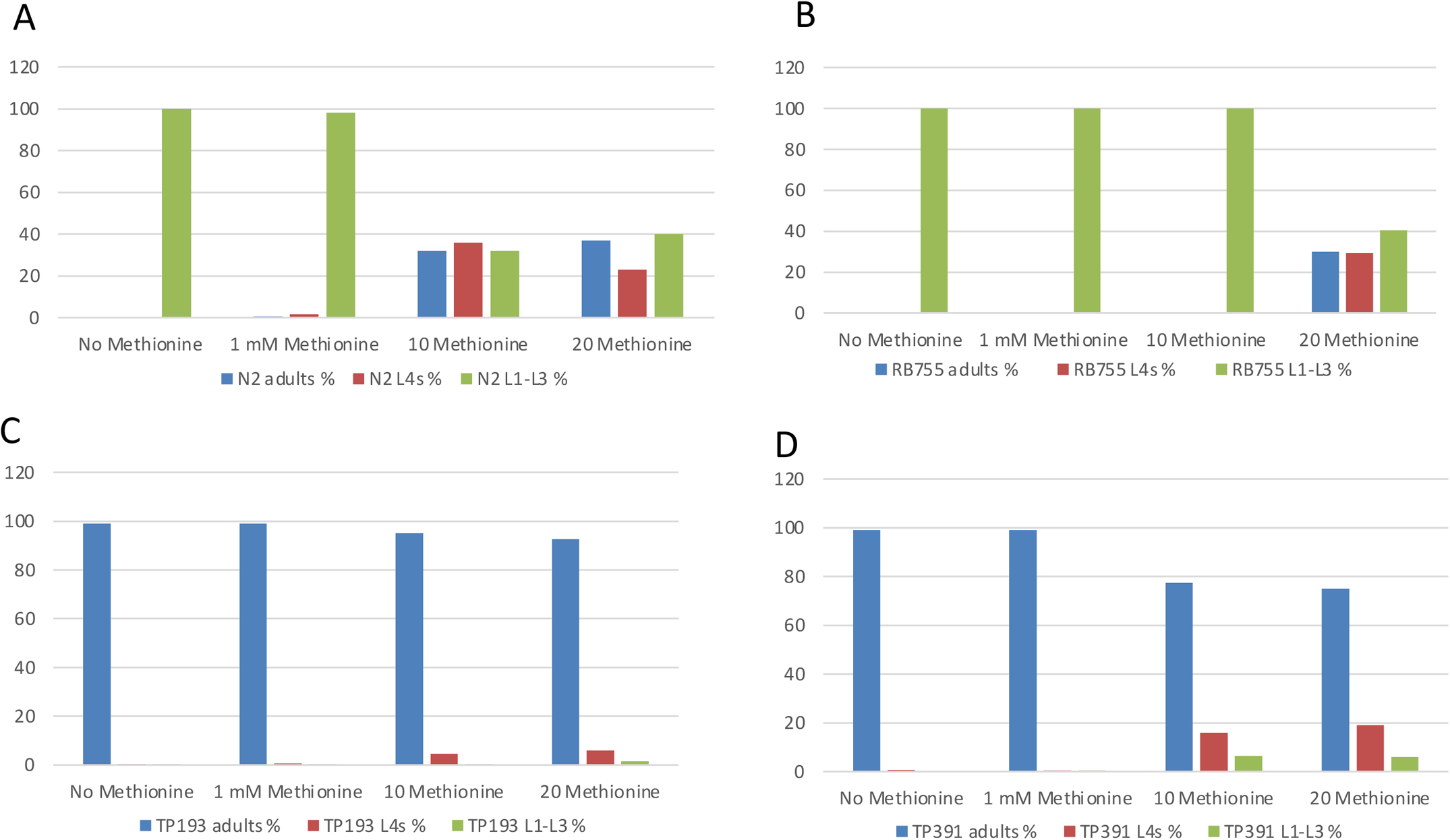
Methionine supplementation partially overcomes DTT-sensitivity in wild type (N2) and *metr-1* mutants. Development (L1 to adult stage over 4 days) of wild type (N2) (A), *metr-1* mutants [RB755, *metr-1*(*ok521*)] (B), *rips-1* mutant [TP193, *rips-1*(*ij109*)] (C), *rips-1*; *metr-1* double mutant mutant [TP391 *metr-1*(*ok521*); *rips-1*(*ij109*)] (D) on 5 mM DTT plates supplemented with 0 mM to 20 mM Methionine. N2 no methionine (n=166), 1 mM methionine (n=259), 10 mM methionine (n=213) and 20 mM methionine (n=119); RB755 no methionine (n=217), 1 mM methionine (n=292), 10 mM methionine (n=140) and 20 mM methionine (n=143); TP193 no methionine (n=244), 1 mM methionine (n=279), 10 mM methionine (n=176) and 20 mM methionine (n=129); TP391 no methionine (n=303), 1 mM methionine (n=259), 10 mM methionine (n=125) and 20 mM methionine (n=89).

### DTT treatment increases *rips-1* transcript abundance

We next used qRT-PCR to assess gene expression changes after exposure of synchronised wild type (N2) L4s to 5 mM DTT for 24 hours, conditions we found to have a minimal effect on worm development within this timeframe. Surprisingly, we found that expression of markers for the endoplasmic reticulum unfolded protein response (*hsp-4*) and mitochondrial unfolded protein response (*hsp-6* and *hsp-60*) were essentially unaffected by this acute exposure to DTT (Figure 8). We also tested two targets of the hypoxia inducible factor (HIF) transcription factor and found transcript abundance for *cysl-2* to be unaffected whereas *nhr-57* transcript was increased ~2-fold with the addition of DTT. We also measured *hsp-4* and *nhr-57* transcript levels in *rips-1* mutant L4s and found similar results to those observed in the wild type genetic background; *hsp-4* mRNA unchanged by DTT and an ~2-fold increase in *nhr-57* mRNA levels. We then tested if DTT affected expression of *rips-1* and found that its abundance was increased ~4.5-fold in wild type treated worms compared to untreated controls (Figure 8). Therefore, in the presence of DTT *C. elegans*, somewhat surprisingly, strongly upregulates expression of *rips-1*, the gene that makes them susceptible to DTT. We also measured *rips-1* transcript levels in an *rips-1* mutant strain and found it was ~6-fold upregulated with DTT treatment. Importantly, this argues against a potential simple explanation for DTT resistance in our mutants, whereby *rips-1* mutation results in loss of DTT uptake.

**Figure 8.**
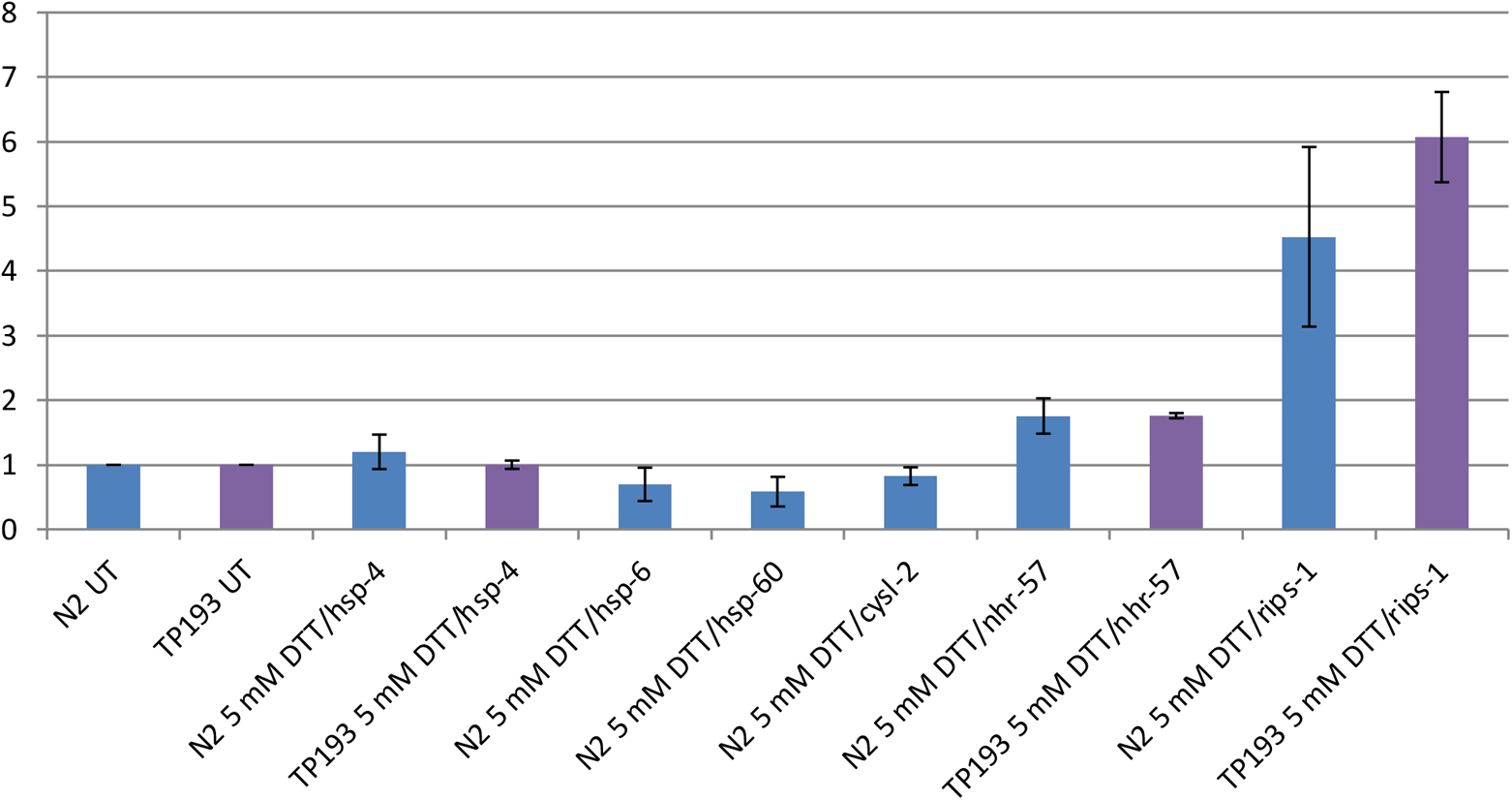
DTT treatment induces *nhr-57* and *rips-1* in wild type and *rips-1* mutant backgrounds. Assessment of gene expression following exposure to 5 mM DTT for 24 hours. Worms were untreated (UT) or exposed to 5 mM DTT, prepped for qRTPCR and the abundance of *hsp-4, hsp-6, cycl-2, nhr-57* and *rips-1* was assessed in both wild type (N2) and a *rips-1* mutant strain [TP193, *rips-1*(*ij109*)].

### RIPS-1 is expressed in the cytoplasm of intestinal and hypodermal cells

To examine regulation and localisation of RIPS-1 we constructed a GFP protein fusion containing the putative *rips-1* promoter sequence, *rips-1* genomic coding sequence fused at its C-terminus to GFP, followed by *rips-1* 3 UTR region. Due to the proximity and proposed function of neighbouring genes (Additional Data Figure 4C) the 3 UTR was experimentally defined as described in Additional Data. The resulting plasmid was microinjected into the syncitial gonad of a *rips-1* mutant strain and transgenic progeny identified by the expression of GFP and the co-injected transgenic *myo-2^prom^*::mCherry pharyngeal marker. Two transgenic strains, TP313 and TP315, were characterised in detail. Both strains carry the transgenic sequences as complex extrachromosomal arrays which are lost naturally in a percentage of their progeny. To ensure that the RIPS-1::GFP fusion transgene was functional we looked at survival of transgene-positive and transgene-negative progeny from these strains, reasoning that if the RIPS-1 in the RIPS-1::GFP transgene was functional then it should return sensitivity to DTT in the *rips-1* mutant background. We found no transgene positive worms developed to L4/adults in the presence of 5 mM DTT compared to 100% of transgene negative *rips-1*mutants (Figure 9A) demonstrating the transgene restores RIPS-1 function. We next used the transgenic strains to examine the tissue-specific expression and sub-cellular localisation of RIPS-1 and found faint but specific transgene expression in the cytoplasm of gut and hypodermal cells (Figure 9B).

**Figure 9.**
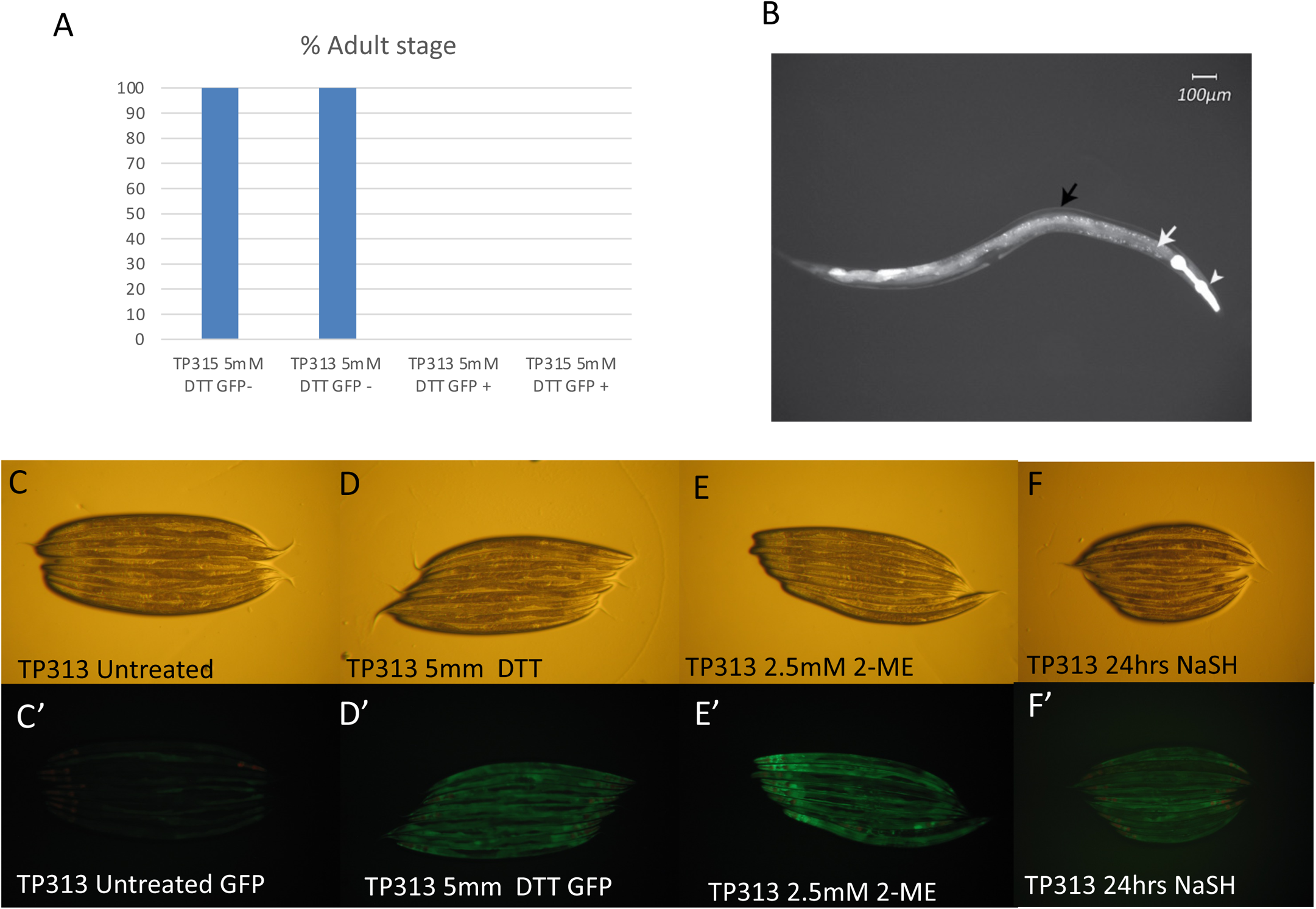
RIPS-1 translational GFP fusion is functionally active, reverses the *rips-1* mutant DTT resistance phenotype, localises to the gut and hypodermis and is induced by thiol reducing agents and hydrogen sulfide. DTT selection of transgenic *rips-1* rescue strain TP313 and TP315 results in selection of GFP positive nematodes that are now sensitive to 5 mM DTT (A). TP315 5 mM DTT GFP- (n=55), TP315 5 mM DTT GFP+ (n=55), TP313 5 mM DTT GFP+ (n=56), TP313 5 mM DTT GFP- (n=56). A translational RIPS-1::GFP fusion (TP313) localises to the hypodermis (black arrow) and the gut (white arrow) with the *myo-2* transformation marker highlighting the pharynx (arrowhead) (B). The RIPS-1::GFP fusion strain TP313 is very weakly expressed under normal culture conditions (C’), but the GFP reporter is induced by 5 mM DTT (D’), 2.5 mM 2-mercaptoethanol (E’) and by sodium hydrogen sulfide (F’) (C-F are the corresponding brightfield images). C-D and C’-D’, images at 25x magnification.

### RIPS-1 is activated by DTT, **β**-mercaptoethanol and H_2_S

We also demonstrated that the RIPS-1::GFP transgene responds to DTT treatment as addition of 5 mM DTT for 24 hours induced strong marker expression in the intestine and hypodermal tissues (Figure 9D and Additional Data Figure 5). Importantly, this verifies our qPCR data, showing that the increase in *rips-1* transcript in response to DTT is also found at the protein level. As *rips-1* mutants are also resistant to 2.5 mM β-mercaptoethanol we tested the effect of this thiol reductant on the marker and found similar upregulation of expression to that observed using DTT (Figure 9D and 9E). We also demonstrated induction of the RIPS-1::GFP marker using DTT with heat-killed *E. coli* OP50-1 and in the absence of bacteria (Additional Data Figure 6). Therefore, as two different thiol reductants upregulate the methyl transferase and as the effect does not require active growing bacteria, we propose that the effect of DTT on RIPS-1 is unlikely to result from a DTT breakdown product or from bacterial metabolism of the compound. Two paralogs of *rips-1*, R08E5.1 and R08F11.4, have been reported as upregulated in the transcriptional response to H_2_S [24]. We therefore tested if this regulation extended to RIPS-1 and found exposure to H_2_S for 24 hours results in moderate RIPS-1::GFP marker induction (Figure 9F and Additional Data Figure 9C).

### Hypoxia inducible factor and mitochondrial dysfunction alter RIPS-1 expression

We next analysed published genome-wide expression-profiling data to identify other conditions and pathways affecting RIPS-1 expression, focussing on studies where *rips-1* was among the final defined gene-sets (as described in more detail in the discussion). *rips-1* transcript abundance has been reported as significantly down-regulated in a SKN-1 gain-of-function mutation that affects mitochondrial function [25], by growth in low selenium [26], and with bacterial food containing high levels of vitamin B12 [27]. *rips-1* transcript abundance was reported as significantly upregulated due to HIF activation [28, 29] and mitochondrial dysfunction [30, 31]. We therefore used our RIPS-1::GFP marker to determine if the results of these genome-wide transcript abundance studies could be replicated at the protein level in a single-gene experiment. We assessed the effect on our RIPS-1::GFP marker of RNAi of the following genes*: vhl-1, rhy-1* and *egl-9*, which result in activation of HIF-1; *clk-1*, *isp-1*, *cyc-1*, and *gas-1* which affect mitochondrial respiration or ubiquinone production [30, 32] and *spg-7*, a mitochondrial protease required for electron transport chain quality control and mitochondrial ribosome biogenesis [31]. Mutation of *clk-1, isp-1*, [33], RNAi of *cyc-1* [34], and RNAi of *spg-7* [31] have all also been shown to activate mitochondrial UPR stress. We also included RNAi of *skn-1* and *mxl-3*. As the *rips-1* transcript was down-regulated in a *skn-1* gain-of-function mutant [25], we predicted that knockdown of *skn-1* by RNAi might produce the opposite effect and result in RIPS-1::GFP marker activation. RNAi of the transcription factor *mxl-3* was included, as deletion of this gene has been shown to reverse the decreased expression of *rips-1* in the *skn-1* gain-of-function mutant background [25]. We did not test the effect of selenium on our RIPS-1 reporter as its low basal expression makes a reduction in expression difficult to demonstrate. RNAi of the mitochondria stress markers *clk-1 and isp-1* failed to induce the RIPS-1::GFP marker (data not shown). However mild induction was observed following RNAi with the mitochondrial stress markers *gas-1* and *cyc-1*. Upregulation of the RIPS-1::GFP marker was also noted in adults following *mxl-3* RNAi and in embryos with *skn-1* RNAi (Figure 10). With the two generation RNAi feeding protocol used we found *spg-7* RNAi resulted in strong developmental arrest phenotype with no detectable marker activation (results not shown). In contrast, Figure 10 (and Additional Data Figure 7) shows that RNAi of *vhl-1, rhy-1* and *egl-9* produce a striking tissue-specific upregulation of the RIPS-1::GFP marker. As RNAi of *vhl-1, rhy-1* and *egl-9* result in HIF-1 activation we then tested if up-regulation of RIPS-1 by DTT was dependant on the HIF-1 signalling cascade. We treated the marker strain with RNAi alone for 3 days then transferred to RNAi with DTT for 24 hours. The empty vector RNAi control and *vhl-1* RNAi both show clear RIPS-1::GFP marker induction with DTT exposure, whereas *hif-1* RNAi strongly suppresses marker induction by DTT (Figure 10 J-N and Additional Data Figure 8).

**Figure 10.**
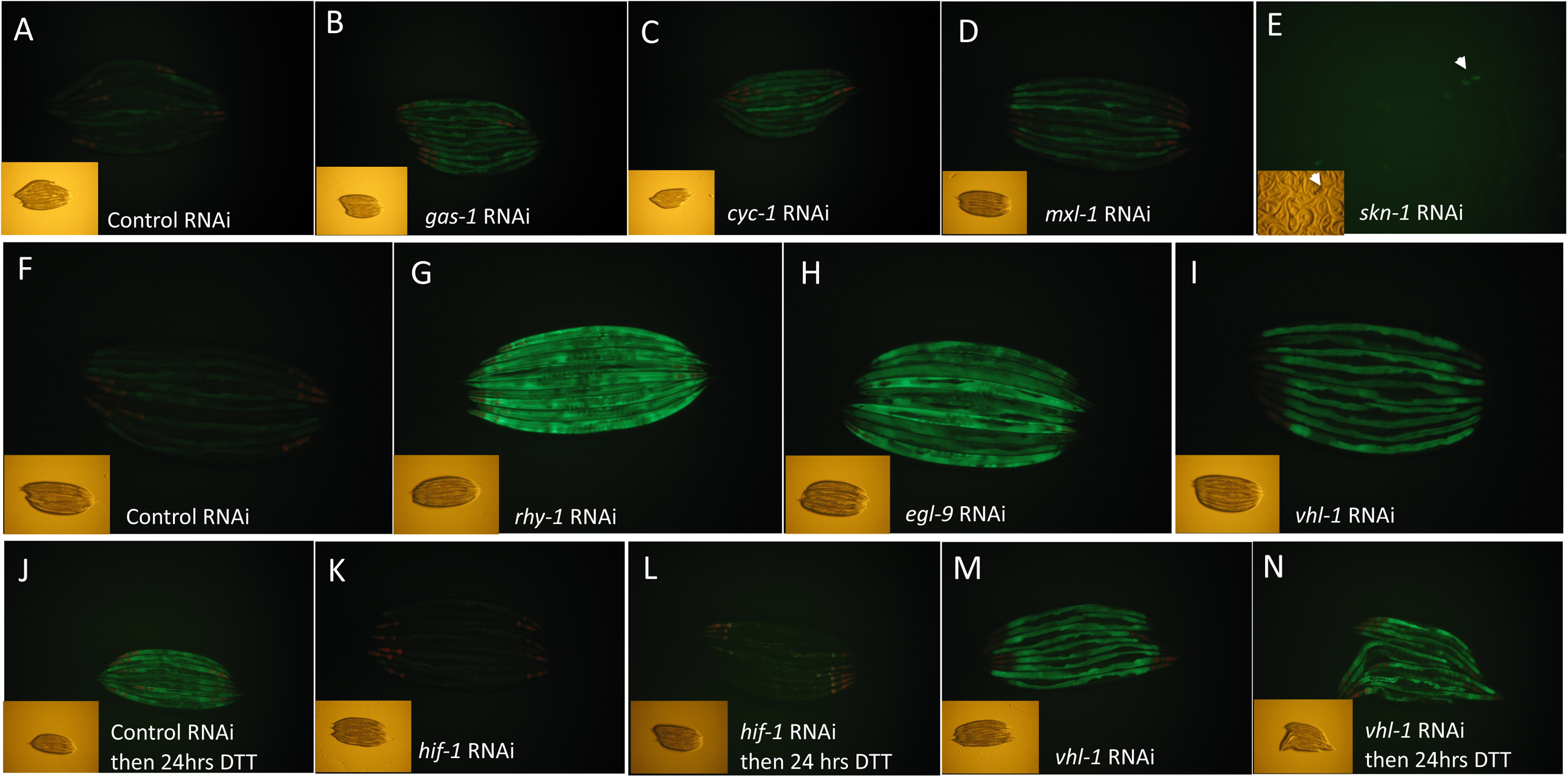
RIPS-1 protein abundance is mildy upregulated by RNAi of mitochondrial electron transport, transcription regulation genes and strongly induced by RNAi of the hypoxia induction pathway genes. Control RNAi causes minimal induction of RIPS-1::GFP (A & F), mild gut and hypodermal induction by RNAi of the mitochondrial electron transport genes *gas-1* (B) and *cyc-1* (C), mild gut-specific induction by the transcription regulator *mxl-1* (D) and embryo-specific induction by the transcription regulator *skn-*1 (E, arrow head). Strong gut and intestine induction was observed following RNAi with the hypoxia pathway associated genes *rhy-1* (G), *egl-9* (H) and induced in a gut-specific manner following *vhl-1* RNAi (I). The transcription factor HIF-1 was shown to control RIPS-1 induction; Control RNAi and 5 mM DTT treatment gave strong RIPS-1::GFP induction (J), *hif-1* RNAi alone failed to induce RIPS-1::GFP (K), *hif-1* RNAi followed by 5 mM DTT treatment also failed to induce RIPS-1::GFP (L). The hypoxia pathway gene *vhl-1* RNAi induced RIPS-1::GFP in the gut tissues (I & M) and this strong induction persisted following 5 mM DTT treatment (N). Bright field images are shown inset; A-D, and J-M magnification x20, insets x5; F-I x25 and insets x8.

**The propionate reporter** *acdh-1* (**VL749) confirms the dietary impact of B12 deficiency** When the mitochondrial *mmcm-1* B12 dependant pathway is disrupted *acdh-1* is induced with the resulting propionate flux in the mitochondria. An *acdh-1* reporter strain has therefore been found to be an excellent marker for B12 deficiency [4]. We used this marker strain to examine the effects of peptone media sources, *E. coli* strains food-sources, and the effect of mitochondrial gene and B12-dependant pathway gene disruption. The marker remains active when normal animal based, but particularly soy-based peptone, is used in NGM plates and when OP50-1 is used as the food source (Figure 11 A). We did however note that this marker is only mildly repressed by the addition of 6.4 nM vitamin B12 (Figure 11B) to the NGM plates but completely repressed by using *E. coli* HT115(DE3) as a food-source in combination with 6.4 nM B12 (Compare Figure 11C to Fig 11D). As expected, RNAi disruption of B12 dependant genes in the cytosol (*metr-1*) and the mitochondria (*mmcm-1*) also induced the marker strain even in the presence of 6.4 nM B12 (Figure 11J and N), but RNAi of neither *rips-1* or *mce-1* affected the induction of this marker in the presence of 6.4 nM B12 (Fig 11 H & L). DTT treatment alone did not affect the induction of this marker (data not shown). The RNAi of genes involved in mitochondrial function, such as *gas-1* (Fig 11 O), *cyc-1* (Fig 11P), and *clk-1* (Fig 11Q), induced this marker even in the presence of 6.4 nM B12 whereas activation of marker following RNAi of rhodoquinone biosynthetic enzyme *kynu-1* was repressed by 6.4 nM B12 (Fig 11R).

**Figure 11.**
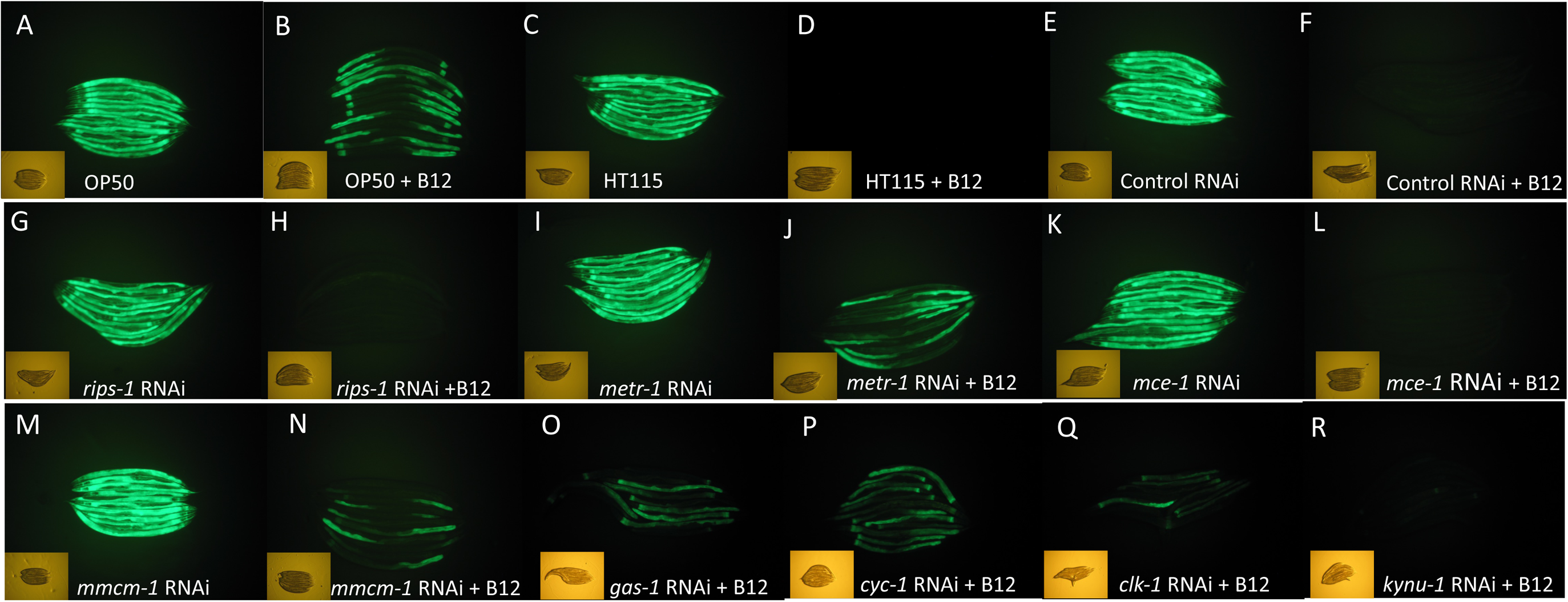
The propionate reporter *acdh-1^prom^*::GFP is induced by diet, RNAi of B12 and mitochondrial pathway genes and is regulated by B12 availability. Worms fed on OP50-1 strongly induce *acdh-1^prom^*::GFP (strain VL749) (A), this induction is only partially suppressed by adding 6.4 nM B12 (B). The RNAi strain HT115(DE3) also induces *acdh-1^prom^*::GFP (C) but this induction is completely suppressed by supplementing with 6.4 nM vitamin B12 (D). Control RNAi [HT115(DE3) and L4440 empty vector] induces *acdh-1^prom^*::GFP (E), but this induction is suppressed by adding 6.4 nM vitamin B12 (F). *rips-1* RNAi induces *acdh-1^prom^*::GFP (G), and induction is suppressed by 6.4 nM B12 (H), *metr-1* RNAi induces *acdh-1^prom^*::GFP (I), and induction is only minimally suppressed by 6.4 nM B12 addition (J), *mce-1* RNAi induces *acdh-1^prom^*::GFP (K), and is completely suppressed by the addition of 6.4 nM B12 (L), *mmcm-1* RNAi induces *acdh-1^prom^*::GFP (M), and induction is minimally suppressed by 6.4 nM B12 (N). RNAi and addition of 6.4 nM B12 only minimally suppresses the induction by *gas-1* RNAi (O), *cyc-1* RNAi (P), *clk-1* RNAi (Q) but 6.4 nM B12 fully suppresses the *kynu-1* RNAi induction of this reporter (R). GFP images shown are at 20x magnification and respective inset brightfield images are at 5x magnification.

## DISCUSSION

### Novel SAM methyl transferase RIPS-1 is involved in methionine dependant sensitivity to thiol reducing agent toxicity

Following several large scale forward genetic screens in *C. elegans* a single novel SAM methyl transferase RIPS-1 (R08E5.3) was identified as conferring resistance to the thiol reducing agent dithiothreitol. We found that that RNAi knock-down of *rips-1* and the presumed loss-of function stop and splice site mutations behaved identically to the amino acid substitutions, indicating that all of the alleles identified for *rips-1* from our mutagenesis screen are loss-of function. It is remarkable that a single methyl transferase enzyme was identified multiple times during separate screens for resistance to thiol reducing agents, represented by a wide range of distinct mutations, nevertheless, the main function of this novel methyl transferase remains to be established.

It is intriguing to speculate that this protein may play a role in adaptation to low oxygen levels, as it is regulated by HIF-1 and RIPS-1 homologues are present in organisms that can adapt to fluctuating O_2_ levels such as marine invertebrates and nematodes. The RIPS-1 methyl transferase is induced by thiol reducing agents and H_2_S, with induction resulting in methionine deficiency that ultimately leads to toxicity; most notably a dramatic developmental delay and ultimately larval lethality. Both these features can be overcome by supplementing with vitamin B12 or methionine. We observe a strong link to the methionine cycle, and via B12-dependance the mitochondrial electron transport chain. Bacterial food sources influences thiol toxicity in *C. elegans* since diet is the sole source of vitamin B12. Therefore, bacterial strains that make sufficient B12 and/or media that contains high levels of B12 allow survival of nematodes in the presence of strong thiol reducing agents. Vitamin B12 has previously been reported to play key roles in *C. elegans* development [3] and availability of this essential vitamin controls the ability of *C. elegans* to break down toxic propionate and imparts metabolic plasticity [4].

### *rips-1* regulation

The SAM methyltransferase *rips-1* was highlighted to be significantly regulated in several genome-wide studies but to date this methyl transferase remains uncharacterised. Two studies also reported that activation of hypoxia inducible factor (HIF-1) resulted in a significant increase in *rips-1* transcript abundance. One study examined mutants in negative regulators of HIF-1 using RNA-seq to profile gene expression regulated by the hypoxia-response pathway under normoxic conditions. *rips-1* was found to be upregulated in each of the three loss-of-function single mutants *egl-9(sa307)*, *rhy-1(ok1402), vhl-1(ok161)* compared to wild type [29]. *rips-1* was moderately down regulated in the HIF single mutant, *hif-1(ia4)*, and in an *egl-9;hif-1* double mutant [29]. Similarly, unpublished work from the Powell-Coffman lab [28] used microarrays to profile expression in *rhy-1(ok1402)* and *egl-9(sa307)* single mutants, and a *swan-1(ok267);vhl-1(ok161)* double mutants under normoxia. They defined a set of 219 genes upregulated in each mutant, with *rips-1* upregulated from 9.8-14.7 fold. Additionally, the authors provided evidence that *rips-1* could be a direct target of HIF-1 direct targets by ChIP-seq.

Two studies have indicated that *rips-1* was changed upon induction of mitochondrial stress. The first of these used microarray of *clk-1(qm30*), *isp-1(qm150)*, and *cyc-1(RNAi)* mutants, which affect mitochondrial respiration or ubiquinone production, to define a set of 73 genes where expression was altered in the same directions in each of the three mutants, with *rips-1* among the most significantly upregulated genes found in the overlap set [30]. The authors showed no effect on lifespan due to knockdown of *rips-1* by RNAi [30], a finding we replicated using a *rips-1* mutant (results not shown). Both *clk-1(qm30)* and *isp-1(qm150)* mutations have been shown to activate the mitochondrial UPR marker *hsp-60^prom^*::GFP [33], while *cyc-1* (C54G4.8) RNAi activates the hsp-*60^prom^*::GFP and *hsp-6^prom^*::GFP mitochondrial UPR markers [34]. The second study used microarrays analysis to identify a set of 685 genes up-regulated due to mitochondrial stress resulting from *spg-7* RNAi [31]. *spg-7* encodes a mitochondrial protease required for electron transport chain quality control and mitochondrial ribosome biogenesis. *rips-1* is induced ~7-fold, making it the ~60^th^ most induced gene in the set, two of the three *rips-1* paralogs were also described in the upregulated transcript set, *R08F11.4* (induced ~43 fold - 5^th^ most induced) and *R08E5.1* (induced ~2.3 fold (~350^th^ most induced) [31].

Profiling of the transcriptional response to H_2_S identified the *rips-1* paralog, R08E5.1, in a set of 17 genes significantly changed by exposure to 50 ppm H_2_S for 1 hour [24]. Additionally, *R08E5.1* and another *rips-1* paralog, *R08F11.4*, were in a set of 30 genes with the greatest increase in transcript abundance after 12 hours exposure to 50 ppm H_2_S. Although *rips-1* was not found in the study we reasoned that, due to regulation of its paralogs by H_2_S and the similarities in DTT, β-mercaptoethanol and H_2_S, *rips-1* may also respond to H_2_S. Exposure of our marker strains to sodium hydrogen sulfide (NaSH) for 24 hrs increased RIPS-1::GFP marker expression. Changes of transcript abundance to H_2_S were also reported to vary dependant on bacterial diet [24] an observation that could potentially be linked to availability of vitamin B12.

A set of 87 core diet-response genes that change depending on the bacterial food source were defined by microarray expression profiling [27]. *rips-1* shows an almost 3-fold downregulation in young adult worms fed with either *E. coli* HT115 or *Comamonas aquatica* DA1877 compared to *E. coli* OP50. The same group also showed that vitamin B12 was the major compound that differed between these strains, with levels of vitamin B12 available to worms much greater in *Comamonas* DA1877 (which synthesises vitamin B12) than in *E. coli* 0P50 [7]. Based on expression of the *acdh-1*::GFP marker the bacterial strain HT115 is thought to provide moderate levels of B12 to the worms [8, 27].

## Conclusions

A proposed model for the role of RIPS-1 in B12 dependant thiol toxicity is depicted in figure 12. Vitamin B12 affects the methionine/SAM pathway having a positive impact on fertility and development, secondly it lowers levels of propionyl-CoA thereby mitigating the toxicity associated with propionic acid build up. When vitamin B12 is limiting, it is conceivable that MMCM-1 and METR-1 compete for this cofactor. As a result, MMCM-1 might be able to utilize more of the vitamin B12 pool for the breakdown of propionic acid when METR-1 is absent. The mechanism by which the *rips-1;mce-1* double mutant is able to enhance resistance to DTT could also be due to the resulting increased B12 availability that can now be used by METR-1 to make methionine. It is interesting to note that the *rips-1*;*metr-1* double mutant survives despite lacking a functioning METR-1, perhaps due to a requirement for balanced levels of methionine or SAM availability in the presence or absence of RIPS-1. In the presence of RIPS-1, methionine/SAM is depleted due to increased RIPS-1 activity, resulting in a decreased survival on DTT whereas, in the absence of RIPS-1, methionine/SAM levels are normal, allowing survival on DTT. In the absence of METR-1, methionine/SAM level is depleted again, also decreasing survival in DTT. In the absence of both RIPS-1 and METR-1, methionine/SAM level are again depleted but not as much as when RIPS-1 is activated in the presence of DTT. This hypothesis is supported by the observation that methionine supplementation also reduced survival of a *rips-1;metr-1* double mutant compared to a *rips-1* single mutant (Fig 7D).

**Figure 12.**
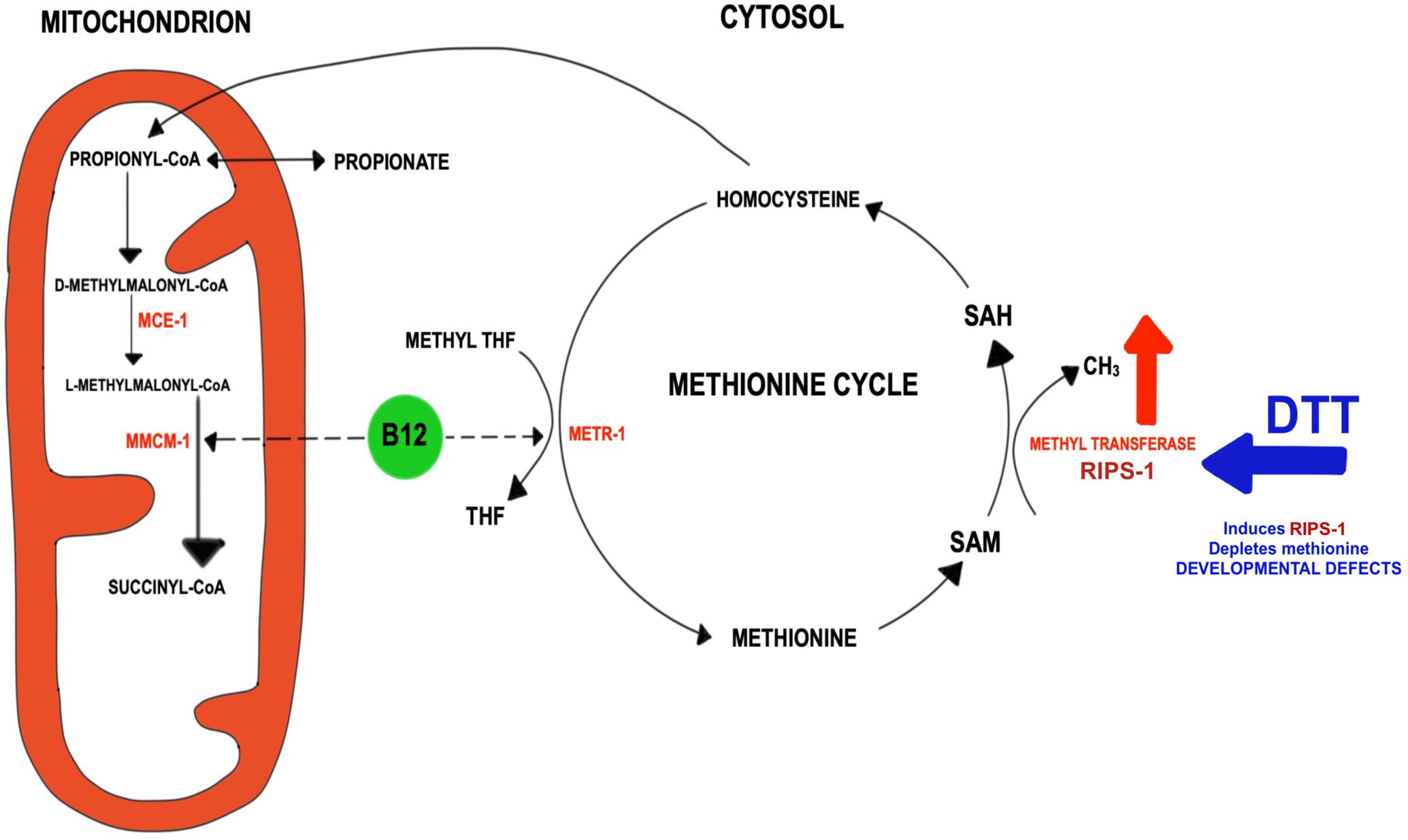
Model for Role of RIPS-1 methyl transferase in the mitochondrial and cytosolic vitamin B12 dependant pathways. Adenosyl Cobalamin (B12) is required in the conversion of TCA cycle dependant Propionyl-CoA to Succinyl-CoA. This process includes the isomerisation of D-methylmalonyl-CoA to L-methylmalonyl-CoA and is catalysed by the enzyme Methylmalonyl-CoA Epimerase (MCE-1), followed by the B12 dependant Methyl Malonyl-CoA Mutase (MMCM-1) conversion to succinyl-CoA, that feeds directly into the TCA cycle. In B12 depletion or disruption of MCE-1 or MMCM-1 this pathway is interrupted and propionyl-CoA is converted into toxic propionate (and the *acdh-1^prom^*::GFP marker is induced). Methyl cobalamin (vitamin B12) is also required for the folate and methionine cycles in the cytosol by acting as an essential co-factor for the MEthyltetrahydrofolate-homocysteine methylTRansferase (methionine synthetase) enzyme (METR-1). This enzyme demethylates methyl tetrahydrofolate (methyl THF) to tetrahydofolate (THF) in the folate cycle and methylates homocysteine to methionine in the methionine cycle. Methionine can be further converted to S-Adenosyl Methionine (SAM), then gets demethylated to S-Adenosyl Homocysteine (SAH). This demethylation reaction is a major source of methyl groups that is catalysed by S-Adenosyl Methionine Methyl Transferases including RIPS-1. The final step of the methionine cycle involves conversion of SAH to homocysteine, which can ultimately feed into the TCA cycle via cystathionine, alpha-ketobutyrate and propionyl-CoA. Thiol reducing agents, including DTT, induce the S-Adenosyl Methionine Methyl Transferase RIPS-1, in turn depleting the available methionine pool. The vitamin B12 dependant reactions including methionine synthetase become rate-limiting and in the absence of B12 supplementation, thiol reducing agents cause developmental defects and lethality.

In nematodes and a limited subset of metazoans including annelids, molluscs, and trematodes, there is an ability to switch from aerobic to anaerobic metabolism by reversing the TCA cycle [36]. This results in reversing the succinate to fumarate conversion by using the unique and highly reduced electron acceptor rhodoquinone. This process also reverses the MMCM-1 dependant cycle thereby removing dependence on adenosyl cobalamin and leads to the production of toxic propionate. It could be hypothesized that one potential function of the HIF-1 induced SAM methyl transferase RIPS-1 is in the biosynthesis of reduced rhodoquinone as an adaptation to anaerobic conditions. Interestingly, RIPS-1 is also listed as containing a ubiquinone domain Ubie_Methyltran (Phenobank; https://worm.mpi-cbg.de/phenobank/cgi-bin/ProteinPage.py?GeneID=511989). The fine balance between this ubiquinone to rhodoquinone switch, reversal of the B12 dependant TCA cycle, and control of propionate build up must be carefully regulated and is highly dependent on the redox poise of the cytosol and the mitochondria. A role for mitochondrial activation was also recognised using an RNAi based approach, where key mitochondrial gene disruption led to *acdh-1* (VL749 marker) propionate induction, this was significant for *gas-1*, *cyc-1* and to a lesser extent *clk-1*.

From our studies, it is clear that RIPS-1 plays a key role in the response to environmental conditions such as H_2_S, O_2_, methionine availability and dietary B12 availability. Intriguingly this uncharacterised SAM methyl transferase is induced by thiol reducing agents only to become sensitive to the associated toxicity via the resulting depletion of methionine. This study highlights the important balance between B12 dependant reactions in the cytosol and mitochondrion and the significant role played by diet in these processes.

## Methods

### Growth and maintenance of *C. elegans*

*C. elegans* strains were maintained as described in [37], strains used in this study are shown in Additional Data Table 3. However, as our DTT assays were found to be sensitive to both bacterial food and growth media (see Results), NGM to be used in DTT assays was made using soybean peptone (Sigma 70178, peptone from soybean, enzymatic digest) and high purity agar (Sigma 05038, agar for microbiology) and only freshly grown bacterial cultures, grown overnight the day before use, were used to seed NGM-DTT plates. DTT was made by dissolving DL-dithiothreitol (D0632 Sigma) to 1M in water (Sigma W4502, Nuclease-Free Water, for Molecular Biology) with single use aliquots stored at −20°C and used within 1 month. DTT was added to NGM at 55°C along with the other standard additives [37]. Set plates were dried for 1 hour in a flow hood, seeded with fresh *E. coli*, left in a 20°C incubator in a plastic box with no lid overnight, and then used the following day. Except where noted, the bacterial food used was *E. coli* OP50-1 (from the Caenorhabditis Genetics Center, CGC) which was grown in standard LB broth with Streptomycin (Sigma) from a 12.5 mg/ml stock dissolved in H_2_0, final concentration 12.5 ug/mL. *E. coli* HT115(DE3) (available from the CGC) transformed with an empty vector RNAi was grown in ampicillin at a final concentration of 100 ug/mL. Heat-killed OP50-1 was produced by heating an overnight culture to 60°C for 30 minutes, prior to plate seeding.

### Embryo preparations and synchronous cultures

Mixed stage cultures were grown on NGM-OP50-1 plates until they contained a large population of gravid hermaphrodites and just prior to the food supply being exhausted. Strains were collected and the embryos were isolated essentially as described [37] but with the following changes; worms were lysed in a total volume of 5 mls using 0.25 mL 5N KOH (final concentration 0.25N) and ~0.06-0.5 mL NaOCl (10%, Sigma 1056142500) (volume dependent on the age of the NaOCl solution) and three wash steps each with 12 mL of M9 buffer. To prepare synchronous cultures, the bleached embryos were added to ~18 mL M9 buffer in a 90 mm petri plate and allowed to grow overnight at 20°C. The resulting starved L1s, which are synchronised at the beginning of the first larval stage, were transferred to 15 mL tubes, washed twice with 12 mL of M9 buffer, added to growth plates using low binding pipette tips, then grown until the required developmental stage.

### Plate supplementation

Vitamin B12 (cobalamin) was purchased in its precursor form cyanocob(III)alamin (CNCbl) (Sigma V2876). CNCbl is converted to the physiologically active forms of vitamin B12, primarily, methylcobalamin (cytoplasmic) and adenosylcobalamin (mitochondrial). CNCbl was prepared as a 10 mM stock in water and used at a concentration of 6.4-64 nM in NGM plates. Methyl THF (Sigma M0132) was added to NGM media during cooling to give concentration of 100µM. Hydrogen sulfide (14.3 M, Sigma, 161527) was diluted to 1M immediately before in water (Sigma, W4502) and added to a concentration of 2.5 mM in NGM plates. L-Methionine (Sigma M9625 was added at 1 to 20mM to NGM media during cooling). To assess the effect of survival in various conditions ~75 embryos, prepared by bleaching as above, were added to 6 cm plates and the number that had developed to adults counted 3-4 days later.

### Microscopy

For high powered microscopy, live *C. elegans* were transferred to slides with a 2% agarose/0.065% sodium azide pad in 5 uL M9 buffer, a coverslip was added and sealed with liquid paraffin. Images were captured using an Axioskop 2 microscope (Zeiss) and an AxioCam MRm camera (Zeiss). Low power group images of worms were taken by picking live worms to a ~10 uL 10 mM sodium azide solution on unseeded NGM agar plates. Images were taken using a Canon PowerShot G6 just as the azide pool dried before worm recovery.

### Ethyl methanesulfonate (EMS) mutagenesis and DTT selection

Synchronised L4 larvae were exposed to freshly diluted 50 mM EMS (Sigma M0880) for four hours at room temperature (ref: “C. elegans: a practical approach”, chapter “Conventional Genetics” by Jonathan Hodgkin, “Protocol 1-EMS Mutagenesis”, pages 252-253) and selected for increased survival NGM-OP50-1 plates with on 5 mM −7.5 mM DTT. A total of approximately 100,000 genomes were mutagenised in three separate screens with each screen being divided into sub-population to identify independent mutants.

### Whole-genome re-sequencing

Strains were prepared for whole-genome re-sequencing and SNP-based mapping [11] by crossing mutant strains with strain CB4856 (Hawaiian) males. As DTT resistant strains are otherwise wild type in morphology and behaviour, crossed progeny were selected by cloning L4 hermaphrodites from the F1 progeny to standard NGM plates and, after egg laying, testing the F1 mother for heterozygosity of a 2940 bp deletion found in the Hawaiian strain (Left flank: AAACCAACGACTCACTAGAGCGCGTATTTT Right flank: TCATCAAATACTTGCATCAACTCCTGAACG; [38]) using primers oADW0165/0166/0168. From crossed F1 plates ~200 F2 L4s were cloned onto NGM OP50-1 plates containing 5 mM DTT. Each resistant strain was then put through a second round of identical selection. To reduce the burden of background mutations some of the DTT resistance alleles were outcrossed two times to N2 prior to characterisation and sequencing, see Additional Data Table 3.

Next Generation Sequencing (NGS) libraries were prepared and sequenced at Glasgow Polyomics (https://www.polyomics.gla.ac.uk/). Briefly, 200 ng genomic DNA, isolated as described in Additional Data - Methods, was fragmented using a Bioruptor Pico (Diagenode) (5 seconds on, 90 seconds off, for 7 cycles), insert size selected for 550 bp, and whole genome libraries constructed using a TruSeq Nano DNA LT Sample Prep Kit (Illumina). Libraries were quantified using a Qubit dsDNA HS kit (Thermo Fisher) with quality assessed using a 2100 Bioanalyzer (Agilent). Two libraries were pooled for sequencing on an Illumina MiSeq using version 3 sequencing reagents with 2 x 300 cycles. NGS data was analysed at the public instance of the open source, web-based platform Galaxy (usegalaxy.org) [39] using the CloudMap [40] “Hawaiian Variant Mapping with WGS Data” and “Variant Calling Workflows”.

### RNAi by bacterial feeding

RNAi was carried out using dsRNA-producing bacteria as a food source. The method was essentially as described [41] but with NGM plates containing 100ug/ml Ampicillin, 1 mM IPTG, plus or minus 5 mM DTT, and using NGM prepared using soybean peptone and high purity agar (as described above). L4s were washed in M9 in a watch glass before being placed on the primary RNAi plates where they were grown for 24 hours then transferred to a secondary RNAi plate. The F1 progeny from the second RNAi plates was scored. All bacterial RNAi strains were from the Ahringer lab *C. elegans* RNAi library [13] or purchased from SourceBioscience. The *rips-1* RNAi clone was sequenced to confirm the identity of the 997 bp insert. In addition to an empty vector negative control RNAi, dpy-11 (F46E10.9) was included in all RNAi experiments as a positive control.

### RIPS-1 GFP translational fusion

A 3529 bp fragment of *rips-1*, extending from 1078 bp upstream to just downstream of the 3’UTR, as defined by 3 RACE, was amplified from N2 genomic DNA using *Pfu* Ultra II Fusion HS polymerase (Agilent) with primers oADW0256/258. The amplicon was purified using a PCR Purification kit (QIAGEN), A-tailed using GoTaq2 (Promega), and ligated into pCR2.1TOPO (Invitrogen) to generate plasmid pADW0105. A QuikChange XL kit (Agilent) was used for site-directed mutagenesis with plasmid pADW0105 [*rips-1*(+), pCR2.1 TOPO] as template, using primers oADW0272/273 and following the manufactures protocol but with the following adjusted cycling parameters: 95°C 2 mins x 1 cycle; 95°C 1 mins, 60°C 1 min, 68 °C 19 mins x 18 cycles; 68 °C 19 mins x 1 cycle. Site-directed mutagenesis removed the stop codon and introduced an *Xma* I restriction site in the sequence and generated plasmid pLBG003. Site-directed mutagenesis results in a change of RIPS-1 C-terminal residues from QN* to QNPG. GFP was amplified from plasmid pPD95.75 (Addgene) using *PfuUltra* II HS DNA Polymerase (Agilent) and primers oADW0274/0275 and cloned into Sma I/CIAP cut pLBG003 to generate plasmid pLBG007 [*rips-1^prom^*::RIPS-1::GFP::*rips-1* 3 UTR] (referred to as RIPS-1::GFP). The identity of plasmid inserts was confirmed by restriction enzyme digest and sequencing using vector primers M13Rev(−29) and M13Uni(−21), while the RIPS-1::GFP junctions were sequenced using oADW0274 (GFP F) and oADW0254 (3’RACE primer). Transgenic strains were created following standard microinjection protocols [42] using a Zeiss Axiovert microscope. The DTT resistant strain TP193 [*rips-1(ij109)*], was microinjected with a mix containing 10 ng/μL pLBG007 [RIPS-1::GFP], 10 ng/μL pADW021(3) [myo-2^prom^::mCherry] (pharyngeal marker), and 100 ng/μL 1 kb ladder (Invitrogen). Four transgenic strains carrying extrachromosomal were generated with strains TP313 and TP315 characterised in detail.

### qRT-PCR

Material for expression analysis by qRT-PCR was prepared by treating the wild type N2 and the DTT resistant strain TP193 with 5 mM DTT for 24 hours. Cultures were synchronised by bleaching, as described above, and starved L1s plated on standard NGM OP50-1 for 48 at 20°C hours until the L4 stage. L4s were washed from plates using M9 buffer, collected in a 15 ml tube, then centrifuged at 1150 x g for 3 minutes at 20°C. The worm pellet was re-suspended in 500 μL M9 and distributed to standard NGM OP50-1 plates (seeded the day before with freshly grown OP50-1) containing either no DTT or 5 mM DTT. After 24 hours the young adults were collected by gentle addition of M9 buffer so that embryos remained in OP50-1 bacterial lawn. Tubes were centrifuged as before, washed once with 12 mL M9 buffer, re-centrifuged, the supernatant removed to 1 mL, transferred to 1.5 ml tube using a low binding tip, centrifuged for 3 mins at 2000 rpm, the supernatant removed to minimum volume, and worms frozen at −80°C. This process was repeated on three separate occasions to generate biological triplicates for qRT-PCR analysis.

Total RNA was isolated, as described in Additional Data - Methods, and 500 ng of DNAse I-treated total RNA used in enzyme-plus and enzyme-minus reverse transcriptions using the AffinityScript qPCR cDNA Synthesis Kit (Agilent) with oligo(dT) primer. Reverse-transcription reactions were diluted with (20 μL + 280 μL H_2_0) and qPCR performed using an Agilent Mx3005P qPCR System following the Brilliant III Ultra-Fast SYBR Green QPCR Master Mix (Agilent Technologies) protocol. Tubulin gamma *tbg-1* (F58A4.8) was used as a normalising gene in all cases with some results confirmed using a second normalising gene, actin *act-3* (T04C12.4). All results are from biological triplicate samples and with each PCR in triplicate.

## Supporting information

Manuscript

## Abbreviations

DTT: dithiothreitol
MMCM: methylmalonyl coenzyme A
MS: methionine synthetase
MCE: methylmalonyl-CoA epimerase
ME: β-mercaptoethanol
H_2_S: hydrogen sulfide
NGM: nematode growth media
SNP: single nucleotide polymorphism.

## ACKNOWLEDGMENTS

Gillian McCormack, Gillian Stepek, Graham Hamilton and Ewan Calder are thanked for helping with DTT selection and WGS. Don Moerman and Mark Edgley are thanked for supplying flanking sequence to allow mapping of a Hawaiian-specific deletion to the updated *C. elegans* genome build. The CGC is thanked for providing many of the *C. elegans* strains used in this study. BBSRC for Funding through the award **BB/K006983/1 to APP.**

## DECLARATIONS

All data described in this manuscript is freely available in the additional data section and will be released to the next update of the public Wormbase database (https://wormbase.org).

All authors declare that they have no competing interests.

## ADDITIONAL INFORMATION

Additional Figures 1 through 8 (**additional figures.pdf**). Contains additional supporting figures.

Additional file 1 (**additional file.xlsx**). Contains detailed Excel files of BLAST comparisons of methyl transferase RIPS-1 (R08E5.3) paralogs and homologs.

Additional Methods and Tables (**Winter B12 Additional Methods and Tables.docx**). Word file that contains detailed methods and supporting data tables.

