## Supplementary material for "Dietary derived Vitamin B12 protects the nematode *Caenorhabditis elegans* from Thiol Reducing agents": Manuscript

**ADDITIONAL DATA**

**Additional Materials and Methods**

**gDNA isolation**

Worms were grown on NGM OP50-1 plates and collected just prior to clearing of the bacterial lawn by washing with M9 buffer and collection in 15 ml tubes followed by centrifugation at 1150 xg for 3 minutes in an AccuSpin centrifuge at 20°C. Worms were washed twice by resuspension in 12 mL M9 followed by centrifugation, re-suspended in M9 and placed on a rocking platform in a 15 ml tube on its side at room temperature for 2 hours, washed twice with M9 and then the worm pellet frozen at -80°C. Genomic DNA was isolated using Gentra Puregene Tissue Kit (QIAGEN) by following the DNA Purification from Tissue protocol but without the liquid nitrogen grinding step. DNA was further purified using a Genomic DNA Clean & Concentrator kit (Zymo Research) and quantified using a Qubit dsDNA Broad Range Assay Kit (Thermo Fisher).

**Sanger sequencing confirmations of *rips-1* alleles in DTT strains**

gDNA was isolated, as described above, from the DTT resistant strains (non-Hawaiian crossed) and a 2429 bp of the *rips-1* locus amplified using *PfuUltra* II (Agilent) with primers oGS0001 and oGS0002**.** PCR products were cleaned using a QIAquick PCR Purification Kit (QIAGEN) and Sanger sequencing using primers oGS0001, oGS0002, oGS0003 and oGS0004. All Sanger sequencing in this study was performed by Eurofins Genomics.

**Total RNA extraction**

Worm pellets frozen at 80°C were re-suspended in 1 mL of TRIZOL Reagent (Invitrogen, ThermoFisher), added to a liquid nitrogen-cooled mortar and pestle then ground to a fine powder. Once thawed, samples were transferred to 1.5 ml tubes and frozen at 80°C. To isolate total RNA, samples were thawed, vortexed for 15 seconds, incubated for 10 minutes at room temperature with occasional vortexing, then centrifuged for 10 minutes at 12,000g at 4°C to pellet the insoluble material. Samples were transferred to 2 ml RNAse-free tubes (Ambion) and a Direct-zol RNA MiniPrep (Zymo Research) protocol with on-column DNase I treatment followed. Purified total RNA was quantified using a QUBIT RNA High Sensitivity Assay kit (Invitrogen, ThermoFisher) and RNA integrity assessed by agarose gel electrophoresis.

**3’RACE for *rips-1***

RNA was isolated from mixed stage and synchronised L4 cultures of wild type N2 as described above. A 3´RACE System for Rapid Amplification of cDNA Ends (Invitrogen, ThermoFisher: 18373019) was used following the manufactures protocol using 1 μg of total RNA from each preparation. PCR was performed on the 3ʹRACE material using GoTaq2 (Promega) with primers AUAP and oADW0254, with secondary amplification using a 1/1000 dilution of primary PCR as template with primers AUAP and oADW0255. PCR products were purified using QIAGEN PCR purification kit, cloned into pCR2.1TOPO and Sanger sequenced with M13Rev(-29), M13Uni(-21).

**Genotyping**

Single worms were lysed in 5 μL of a Proteinase K solution (0.5 μg/μL final concentration proteinase K; added immediately before use to a buffer containing 10 mM Tris (pH 8.0), 50 mM KCl, 2.5 mM MgCl2, 0.45% Tween-20 and 0.05% gelatine) by incubating for 1 hour at 65°C followed by 95°C for 15 minutes and the lysis reaction used directly as PCR template using GoTaq G2 Flexi DNA Polymerase (Promega). The DTT resistant amino acid substitution allele *rips-1 (ij109)* was genotyped by Sanger sequencing. Following genetic crosses and selection on 5 mM DTT, PCR using primers oGS0003 and oGS0002 on single-worm lysates was used to amplify *rips-1* genomic sequence flanking the *ij109* allele with the product purified using a QIAquick PCR Purification Kit (QIAGEN) and then Sanger sequenced using oGS0003 and oADW0279. The *mce-1* (*ok243*) deletion breakpoints were first defined by Sanger sequencing of a PCR product generated using primers mce-1ok243 seq IL and mce-1ok243 seq IR. *rips-1 (ij109)* [strain TP193] was crossed with *mce-1(ok243)* [strain RB512] to create the double mutant strain TP390. The *mce-1* deletion genotype was followed using primers *mce-1F*/*mce-1F2*/*mce-1R*, and with the *rips-1(ij109)* point mutation confirmed as described above. *rips-1 (ij109)* [TP193 strain] was crossed with *metr-1(ok521)* [strain RB755] to create the double mutant strain TP391 with the *metr-1(ok521)* allele being followed using primers oADW0326/ oADW0327/ oADW0328.

**Additional Results**

**3ʹ untranslated region from *rips-1 (R08E5.3)***

We noticed that the *rips-1* transcript is annotated in NCBI (NM_071644.5) and WormBase with 1112 bp 3ʹUTR. *R08E5.2 (cysl-3)* is directly downstream of *rips-1* encoded on the opposite strand. The predicted 3ʹUTR of *rips-1* would therefore overlap significantly with coding regions of this neighbouring gene. To determine if this region of *rips-1* would be required for any expression construct and, due to the predicted functions of *cysl-3* in hydrogen sulphide interaction, if there might be a potential interaction between these two genes at the RNA level, 3ʹRACE was performed to experimentally define the 3ʹUTR for *rips-1*. Sequencing of 3ʹRACE products from mixed developmental stage and stage-specific (synchronised L4s) was analysed which revealed that the position of poly-A addition site for longest product identified is 177 nucleotides after the annotated stop. This results in an overlap region of only 76 bp involving only 12 bp if *cysl-3* coding sequence making any interaction at the RNA level unlikely.

**Additional Tables**

**Additional Table 1. RNAi mini-screen for B12-related pathways**

| **Gene** | **Cosmid ID** | **Description** | **Pathway** |
| --- | --- | --- | --- |
| *cblc-1* | ZK546.17 | Ortholog of human MMACHC (methylmalonic aciduria (cobalamin deficiency) cblC type, with homocystinuria)  CblC. | B12 processing |
| _ | Y76A2B.5 | Ortholog of MMADHC (methylmalonic aciduria and homocystinuria, cblD type)  CblD. | B12 processing |
| *sams-1* | C49F5.1 | S-adenosylmethionine synthase | Methionine |
| *ahcy-1* | K02F2.2 | S-AdenosylhomoCysteine HYdrolase homolog | Methionine |
| *metr-1* | R03D7.1 | Methionine synthetase  CblG | Methionine |
| *mtrr-1* | C01G6.6 | Methionine synthetase reductase  CblE | Methionine |
| *mthf-1* | C06A8.1 | Orthologous to human methylenetetrahydrofolate reductase (MTHFR) | Folate |
| *mel-32* | C05D11.11 | serine hydroxymethyltransferase | Folate |
| *cbs-1* | ZC373.1 | CYSTATHIONINE-BETA-SYNTHASE | Transulpheration |
| *cysl-1* | C17G1.7 | CYsteine Synthase Like | Transulpheration |
| *cysl-2* | K10H10.2 | CYsteine Synthase Like | Transulpheration |
| *cysl-3* | R08E5.2 | CYsteine Synthase Like | Transulpheration |
| *cbl-1* | C12C8.2 | putative cystathionine gamma-lyase orthologous to human CTH | Transulpheration |
| *cth-1* | F22B8.6 | putative cystathionine gamma-lyase; CTH-1 is orthologous to human CTH | Transulpheration |
| *cth-2* | ZK1127.10 | putative cystathionine gamma-lyase; CTH-2 is orthologous to human CTH | Transulpheration |
| *gcs-1* | F37B12.2 | C. elegans ortholog of gamma-glutamine cysteine synthetase heavy chain (GCS(h) | Transulpheration |
| *gss-1* | M176.2 | C. elegans ortholog of glutathione synthetase (GSS) | Transulpheration |
| *mmaa-1* | T02G5.13 | Methylmalonic aciduria type A protein | AdoCbl processing |
| *mmab-1* | C26E6.11 | Methylmalonic aciduria type B protein, co(I)balamin adenosyltransferase (MMAB) | AdoCbl processing |
| *pcca-1* | F27D9.5 | Propionyl-CoA carboxylase alpha chain | Canonical propanoic acid |
| *pccb-1* | F52E4.1 | Propionyl-CoA carboxylase beta chain | Canonical propanoic acid |
| ***mce-1*** | D2030.5 | Methylmalonyl-CoA epimerase | Canonical propanoic acid |
| *mmcm-1* | ZK1058.1 | Methylmalonyl-CoA mutase (MCM) | Mitochondrial B12 requiring enzyme |
| *acdh-1* | C55B7.4 | Acyl CoA DeHydrogenase | Propanoic acid shunt |

**Additional Table 2. DTT resistance alleles**

| **Allele** | **Genomic co-ords (WS220)** | **Nucleotide change**  ***rips-1***  **NM_071644.5** | **Amino acid change**  **RIPS-1**  **NP_504045.1** | **Sequencing (strain used)** |
| --- | --- | --- | --- | --- |
| *ka13* | V: 3772286: G>A | c.49-1 G>A (caG/caA) | Exon 2 splice acceptor | WGS (TP173)  Sanger (TP278) |
| *ka14* | V: 3772286: G>A | c.49-1 G>A (caG/caA) | Exon 2 splice acceptor | WGS (TP174)  Sanger (TP251) |
| *gk902193(^a^)* | V: 3772550: G>A | c.264: tgG/tgA | p.W88* | Sanger (TP295) |
| *ij109* | V: 3773675: T>A | c.400 T>A (Tac/Aac) | p.Y134N | WGS (TP193)  Sanger (TP193) |
| *ka23* | V: 3773814: T>A | c.539 T>A (cTg/cAg) | p.L180Q | WGS (TP183)  Sanger (TP276) |
| *ka29* | V: 3773814: T>A | c.539 T>A (cTg/cAg) | p.L180Q | Sanger (TP189) |
| *ka18* | V: 3773823: G>A | c.548 G>A (gGg/gAg) | p.G183E | Sanger (TP178) |
| *ka28* | V: 3773874: C>T | c.599 C>T (tCg/tTg) | p.S200L | Sanger (TP188) |
| *ka22* | V: 3774050: C>T | c.775 C>T (Cag/Tag) | p.Q259* | Sanger (TP182) |
| *ka9* | V: 3774121: C>T | c.799 C>T (Caa/Taa) | p.Q267*  (p.P116L) | WGS (TP169)  Sanger (TP169) |
| *ka15* | V: 3774133: C>T | c.811 C>T (Cga/Tga) | p.R271*  (p.A150V) | WGS (TP175)  Sanger (TP252) |
| *ka20* | V: 3774245: C>T | c.923 C>T (tCc/tTc) | p.S308F | Sanger (TP180) |
| *ka11* | V: 3774287: C>T | c.965 C>T (gCt/gTt) | p.A322V | WGS (TP171)  Sanger (TP277) |
| *ka21* | V: 3774308: G>A | c.986 G>A (tGg/tAg) | p.W329* | WGS (TP181)  Sanger (TP279) |

^a^ *C. elegans* Million Mutation Project allele.

**Additional Table 3. *C. elegans* strains**

For the DTT resistance alleles ij109 and ka9, outcrossed strains were used both for mapping/WGS and for Sanger sequencing. For six DTT resistance alleles, non-outcrossed strains were used for mapping/WGS and outcrossed strains used for Sanger sequencing. For five alleles only Sanger sequencing was performed, done using non-outcrossed strains. For DTT resistance allele ka17 a possible large re-arrangement was indicated.

| **Strain** | **Genotype** | **Description** | **Source and references** |
| --- | --- | --- | --- |
| TP193 | *rips-1 (ij109)* V | DTT resistant mutant. 0utcrossed twice to N2.  Used for Hawaiian mapping/NGS.  Sanger sequenced for *rips-1*. | This study |
| TP169 | *rips-1 (ka9)* V | DTT resistant mutant. Outcrossed twice to N2.  Used for Hawaiian mapping/NGS.  Sanger sequenced for *rips-1*. | This study |
| TP171 | *rips-1 (ka11)* V | DTT resistant mutant. Not outcrossed.  Used for Hawaiian mapping/NGS. | This study |
| TP277 | *rips-1 (ka11)* V | DTT resistant mutant. Outcrossed twice to N2.  Sanger sequenced for *rips-1*. | This study |
| TP173 | *rips-1 (ka13)* V | DTT resistant mutant. Not outcrossed.  Used for Hawaiian mapping/NGS. | This study |
| TP278 | *rips-1 (ka13)* V | DTT resistant mutant. Outcrossed twice to N2.  Sanger sequenced for *rips-1*. | This study |
| TP174 | *rips-1 (ka14)* V | DTT resistant mutant. Not outcrossed.  Used for Hawaiian mapping/NGS. | This study |
| TP251 | *rips-1 (ka14)* V | DTT resistant mutant. Outcrossed twice to N2.  Sanger sequenced for *rips-1*. | This study |
| TP175 | *rips-1 (ka15)* V | DTT resistant mutant. Not outcrossed.  Used for Hawaiian mapping/NGS. | This study |
| TP252 | *rips-1 (ka15)* V | DTT resistant mutant. Outcrossed twice to N2.  Sanger sequenced for *rips-1*. | This study |
| TP181 | *rips-1 (ka21)* V | DTT resistant mutant. Not outcrossed.  Used for Hawaiian mapping/NGS. | This study |
| TP279 | *rips-1 (ka21)* V | DTT resistant mutant. Outcrossed twice to N2.  Sanger sequenced for *rips-1*. | This study |
| TP183 | *rips-1 (ka23)* V | DTT resistant mutant. Not outcrossed.  Used for Hawaiian mapping/NGS. | This study |
| TP276 | *rips-1 (ka23)* V | DTT resistant mutant. Outcrossed twice to N2.  Sanger sequenced for *rips-1*. | This study |
| TP178 | *rips-1 (ka18)* V | RIPS-1 DTT resistant mutant, not outcrossed.  Sanger sequenced for *rips-1*. | This study |
| TP180 | *rips-1 (ka20)* V | DTT resistant mutant, not outcrossed.  Sanger sequenced for *rips-1*. | This study |
| TP182 | *rips-1 (ka22)* V | DTT resistant mutant. Not outcrossed.  Sanger sequenced for *rips-1*. | This study |
| TP188 | *rips-1 (ka28)* V | DTT resistant mutant. Not outcrossed.  Sanger sequenced for *rips-1*. | This study |
| TP189 | *rips-1 (ka29)* V | DTT resistant mutant. Not outcrossed.  Sanger sequenced for *rips-1.* | This study |
| TP177 | *ka17* | DTT resistant mutant. Not able to outcross.  Used for Hawaiian mapping/NGS. Mapping to chromosome I, possible large re-arrangement. No variant in *rips-1* found by NGS or Sanger sequencing. |  |
| VC40962 | *rips-1* (gk902193) V | *C. elegans* Million Mutation Project strain | Caenorhabditis Genetics Center (CGC) |
| TP295 | *rips-1* (gk902193) V | Outcrossed 4 times to N2, selected on DTT.  Sanger sequenced for *rips-1*. | This study |
| TP313 | *rips-1 (ij109)* V; kaEx [pLBG007 (RIPS-1::GFP) + pADW021(3) (P*myo-2*::mCherry)] | DTT resistant strain carrying RIPS-1::GFP extrachromosomal array | This study |
| TP315 | *rips-1(ij109)* V; kaEx [pLBG007 (RIPS-1::GFP) + pADW021(3) (*Pmyo-2*::mCherry)] | DTT resistant strain carrying RIPS-1::GFP extrachromosomal array | This study |
| VL749 | wwIs24 [*acdh-1p*::GFP + unc-119(+)] | Propionic acid marker | CGC. [1] |
| RB755 | R03D7.1(ok521) II | Methionine synthetase (*metr-1*) mutant | CGC. [2] |
| RB512 | D2030.5(*ok243*) I | Methylmalonyl-CoA Epimerase (*mce-1*) mutant | CGC. [2] |
| TP390 | D2030.5(ok243) I; *rips-1* (ij109) V. | *rips-1* and Methylmalonyl-CoA Epimerase (*mce-1*) double mutant | This study |
| TP391 | *metr-1(ok521)* II; *rips-1* (ij109) V] | *rips-1* and Methionine synthetase (*metr-1*) double mutant | This study |

**Additional Table 4. Oligonucleotide primers**

Sequences are shown 5′-3′. Engineered bases are underlined and restriction enzymes sites are shown in bold font. All oligonucleotide primers were synthesised at Eurofins Genomics. Primers for qPCR and site-directed mutagenesis were HPLC purified.

| **Genotyping primers for a 2940 bp deletion on chromsome V of the Hawaiian strain CB4856** | | |
| --- | --- | --- |
| oADW0165 | GGGATCACCATATTTGGTAAGA (22) | F1 primer, flanks deletion. |
| oADW0166 | CATCGTGATGAAAAGTTGATGAC (23) | R1 primer, flanks deletion. |
| oADW0168 | CCAGTAATGCTTCAGACAAGT (21) | F2 primer, internal to deletion. |

| **Amplification and Sanger sequencing of *rips-1* genomic locus** | | |
| --- | --- | --- |
| oGS0001 | CTACACAACACGTGGACAAC (20) | Amplification forward and sequencing primer. |
| oGS0002 | GTATTCCCCAGCCAGCCATG (20) | Amplification reverse and sequencing primer. |
| oGS0003 | GTAATCGTGAGGTACTCATAC (21) | Sequencing primer. |
| oGS0004 | GTAATGGAACAATCTGACAC (20) | Sequencing primer. |

| **Amplification and Sanger sequencing of allele *rips-1* (*ij109)* for confirmation of genotype after crossing** | | |
| --- | --- | --- |
| oGS0003 | GTAATCGTGAGGTACTCATAC (21) | Amplification forward primer. |
| oGS0002 | GTATTCCCCAGCCAGCCATG (20) | Amplification reverse primer. |
| oADW0279 | GGAGTCCGTCCAGATTTCTG (20) | Sequencing primer. |

| ***rips-1* 3’RACE** | | |
| --- | --- | --- |
| oADW0254 | GGATGTTCGCTATGGTAGAG (20) | Forward primer for 3’RACE. |
| oADW0255 | GGCTCCAGTAACGTATACAC (20) | Nested forward primer for 3’RACE. |
| AUAP | GGCCACGCGTCGACTAGTAC (20) | Abridged universal amplification primer. |

| **RIPS-1::GFP translational fusion** | | |
| --- | --- | --- |
| oADW0256 | GCAGGTAGGCAGGCATAGAA (20) | *rips-1* amplification forward primer. includes a small amount of R08E5.1 coding sequence. |
| oADW0258 | CGATGAGGCGATGGGAATTG (20) | *rips-1* amplification reverse primer. Designed downstream of longest experimental (3’RACE from this study) 3’UTR for *rips-1* and just into 3’ end of *cysl-3* coding sequence |
| oADW0272 | GTATTGTGCCCAGAAAAA**CCC**GGGTACTATTGAAGCC (37) | Site-directed mutagenesis forward primer. Used on *rips-1* rescue construct to mutate stop codon and introduce and an *Xma* I restriction site. |
| oADW0273 | GGCTTCAATAGTACCC**GGG**TTTTTCTGGGCACAATAC (37) | Site-directed mutagenesis reverse primer. |
| oADW0274 | AA**CCCGGG**TATGAGTAAAGGAGAAGAACTTTTCAC (35) | GFP amplification forward primer with *Xma* I restriction site. Additional “T” to adjust reading frame for insertion after R08E5.3. |
| oADW0275 | AA**CCCGGG**CTATTTGTATAGTTCATCCATGCCAT (34) | GFP amplification reverse primer with *Xma* I restriction site |
| oADW0254 | GGATGTTCGCTATGGTAGAG (20) | Designed for 3’RACE, also used to sequence site-directed mutagenized plasmids over the engineered change. |

| **qRT-PCR (one primer of pair was designed over an exon-exon boundary to reduce signal from residual gDNA).** | | |
| --- | --- | --- |
| oADW0306 | AGGCTGCAAAGAGAGGTACA (20) | R08E5.3 F. Designed to be specific for *rips-1* and should not amplify paralog R08E5.1, K12D9.1 or R08F11.4 |
| oADW0307 | TCGTCGAGAGGCTCCAAC (18) | R08E5.3 R. Designed to be specific for *rips-1* and should not amplify paralogs |
| oADW0308 | CACGTGGAGAATGTGAAGGAC (21) | R08E5.1 F. Designed to be specific for R08E5.1 |
| oADW0309 | GCTTCTCGACAACTCCGTGA (20) | R08E5.1 R. Designed to be specific for R08E5.1 |
| oADW0114 | GCCAACGATCAAGGAAACAG (20) | hsp-4 (F43E2.8) F |
| oADW0115 | GATCCAACCTTCACCTCAAC (20) | hsp-4 (F43E2.8) R |
| oADW0312 | GAGGTTCAAAAGGACTTAAAGG (22) | hsp-6 (C37H5.8) F |
| oADW0313 | GTAGCTTGACGCTGAGAATC (20) | hsp-6 (C37H5.8) R |
| oADW0314 | CACTATGGGCCCAAAAGGAA (20) | hsp-60 (Y22D7AL.5) F |
| oADW0315 | CTTGACGAATGCTCTCGAATC (21) | hsp-60 (Y22D7AL.5) R |
| oADW0322 | GCCCACACAAAATTCAAGGCAT (22) | cysl-2 (K10H10.2) F |
| oADW0323 | GTCGCTTGGCCAACTGAAC (19) | cysl-2 (K10H10.2) R |
| oADW0320 | GGCTATCGCTTGCAAAGCAT (20) | nhr-57 (T05B4.2) F |
| oADW0321 | GGCTTGTTGGATTGCTTGAAC (21) | nhr-57 (T05B4.2) R |
| oADW0118 | AAGATCTATTGTTCTACCAGGC (22) | tbg-1 (F58A4.8, tubulin gamma). Primer from [3] |
| oADW0119 | CTTGAACTTCTTGTCCTTGAC (21) | tbg-1 (F58A4.8, tubulin gamma) R. Primer from [3] |
| oADW0310 | GCTCTTGCCCCATCAACC (18) | act-3 (T04C12.4, actin) F |
| oADW0311 | GAGAAAATGGTAAGGCGAAGAG (22) | act-3 (T04C12.4, actin) R |

| **Genotyping *mce-1* and *metr-1* mutant strains** | | |
| --- | --- | --- |
| mce-1ok243 seq IL | CCAAGTAGCCTTCATCTCGC (20) | *mce-1* (D2030.5) allele *ok243*. Primer to define deletion. Primer from [https://cgc.umn.edu/strain/RB512](about:blank) |
| mce-1ok243 seq IR | CCCATGTGCGTAAGGAATTT (20) | *mce-1* (D2030.5) allele *ok243*. Primer to define deletion. Primer from [https://cgc.umn.edu/strain/RB512](about:blank) |
| *mce-1F* | GCAAGTTTACAGCGGGTTCATTG (23) | mce-1 genotyping primer F1 |
| mce-1F2 | AAGCGAAGAATGTCAATAATCAG (23) | mce-1 genotyping primer F2 |
| mce-1R | GAGACTGAACGATGTTGAATAAC (23) | mce-1 genotyping primer R1 |
| oADW0326 | CGAGGATGAAGGAGTTCCAG (20) | *metr-1* (R03D7.1) allele ok521 genotyping F1. Primer from [https://cgc.umn.edu/strain/RB755](about:blank) |
| oADW0327 | GATGAAAGAATCAGACCACTTCG (23) | *metr-1* (R03D7.1) allele ok521 genotyping R1 |
| oADW0328 | TAAACTTGTCCCAGTCGATCTTG (23) | *metr-1* (R03D7.1) allele ok521 genotyping R2 |

**Additional Table 5. Plasmids**

| **Name** | **Description** |
| --- | --- |
| pADW0102(1) | *rips-1* 3’RACE mixed stage, pCR2.1 TOPO |
| pADW0103(12) and (17) | *rips-1* 3’RACE L4, pCR2.1 TOPO |
| pADW0105 | *rips-1* (+), pCR2.1TOPO |
| pLBG003 | *rips-1* (*Xma* I), pCR2.1TOPO |
| pLBG004 | GFP (*Xma* I/*Xma* I), pCR-Blunt II-TOPO |
| pLBG007 | *rips-1* *^prom^*::RIPS-1::GFP::RIPS-1 |
