## Additional figures 1-8 for "Dietary derived Vitamin B12 protects the nematode *Caenorhabditis elegans* from Thiol Reducing agents"

### Additional Figure 1 A

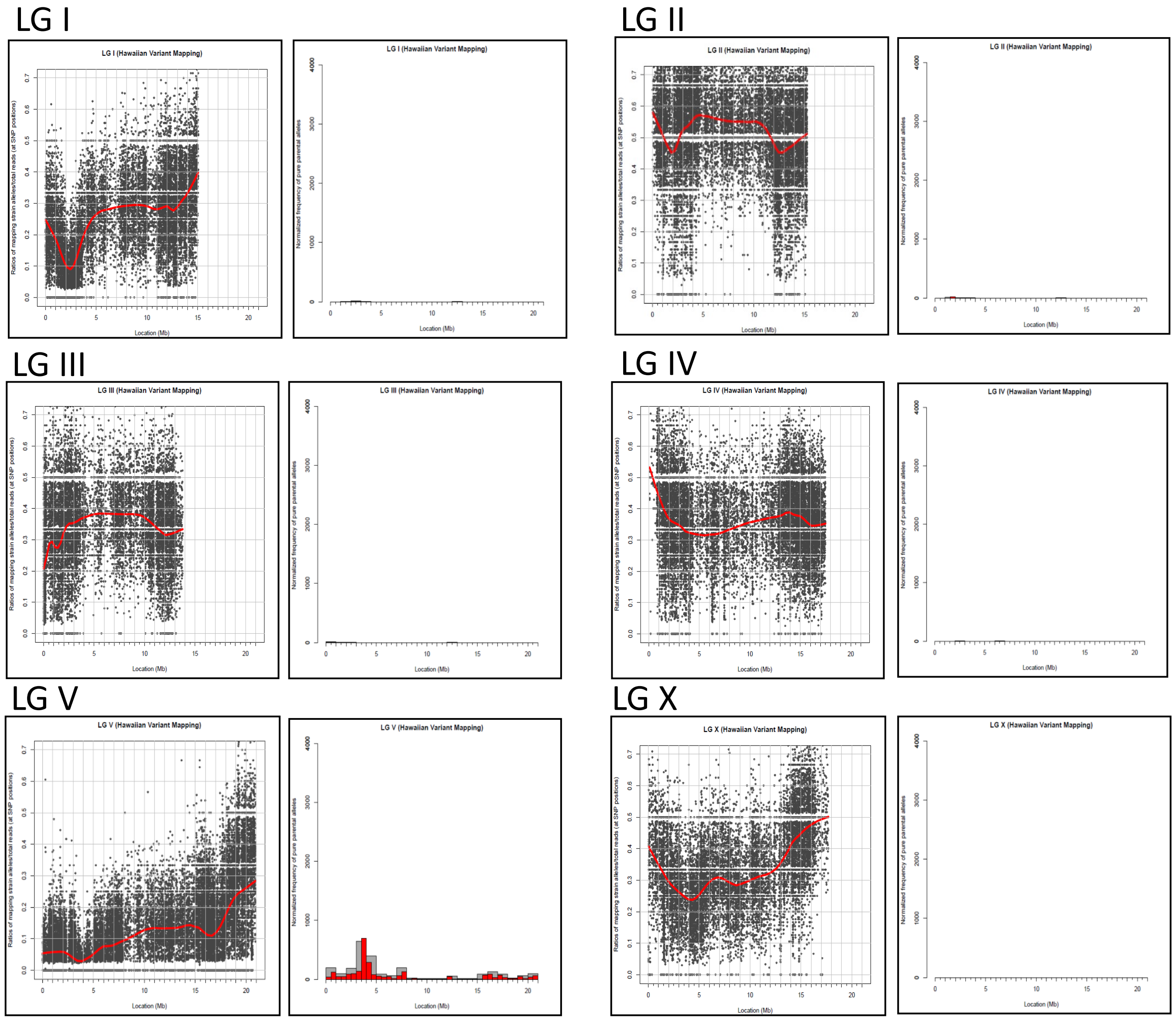


HA mapping output from DTT resistance screen for *rips-1* allele *ij109* (strain TP193). Clear peak visible on Chromosome V.

**Additional Figure 1B**

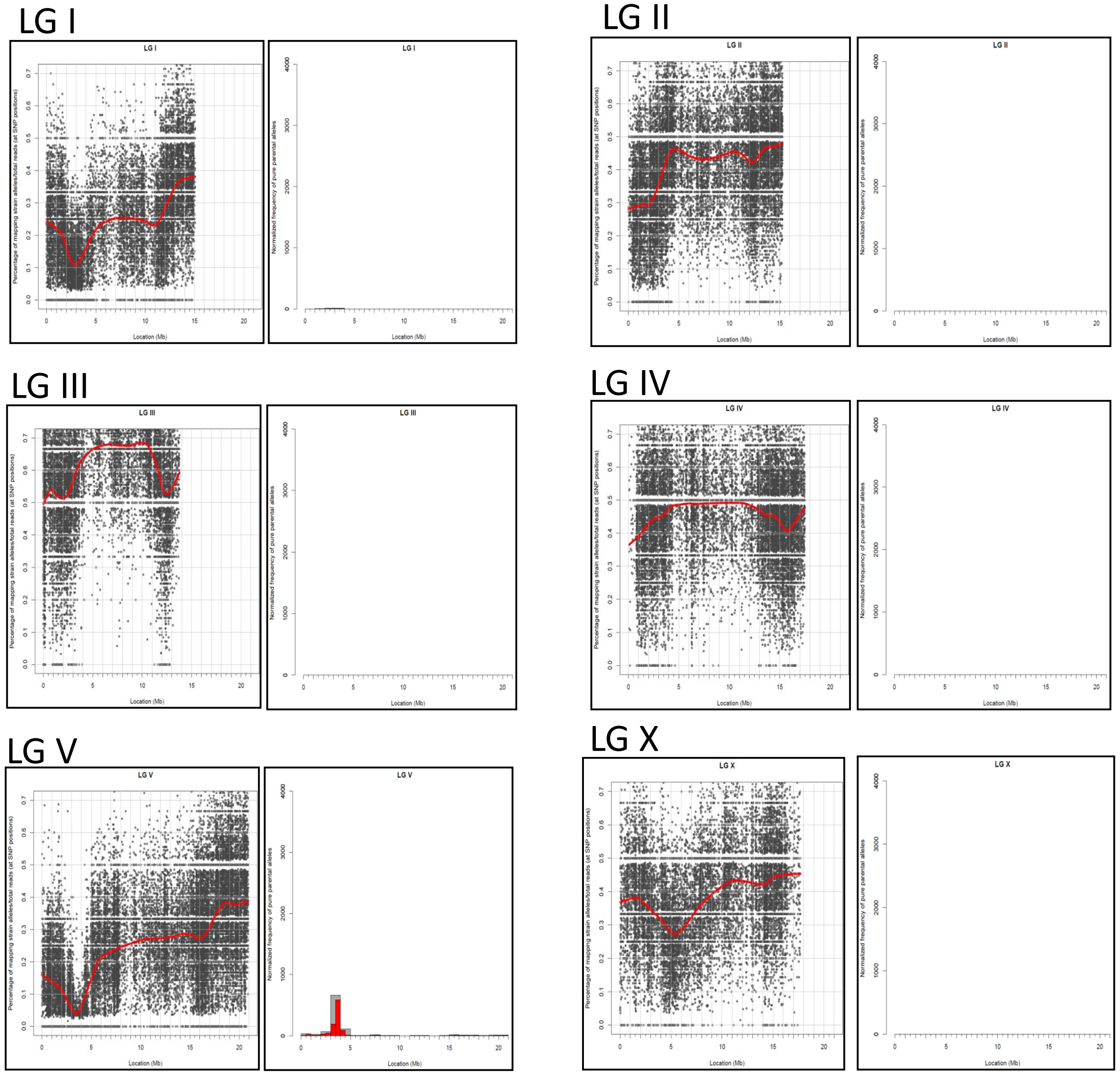


HA mapping output from DTT resistance screen for *rips-1* allele *ka14* (strain TP251). Clear peak visible on Chromosome V.

**Additional Figure 2**

High degree of identity between *rips-1* (R08E5.3) and closest homologue R08E5.1 that will account for potential RNAi cross-reaction.

Regions of ≥20 with 100% nucleotide identity are shown in bold and underlined.
Regions of ≥17 with 100% nucleotide identity are underlined.

Section of R08E5.1 RNAi fragment aligned against unspliced rips-1 (R08E5.3) mRNA

```
              R08E5.1 RNAi frag    (529) GAAGCATGTGATAACTGATATGCTTCCGGACATCGGTCACGGAGTTGTCG
rips-1  unspliced (from WB)   (1691) GAAGCATGTGGTAGCAGATATGCTTCCGGACATCGGCAACGGAGTCGTCG
                        Consensus   (1701) GAAGCATGTG TA C GATATGCTTCCGGACATCGG  ACGGAGT GTCG
                                           1751                                           1800
              R08E5.1 RNAi frag    (579) AGAAGCTTGAAACCGGTGGAATGCGGGTTCTGGACGTCGGGTGTGGTGGT
rips    unspliced (from WB)   (1741) AGAAGCTTGAAGCCGGCGGAGTACGAGTCCTGGACGTCGGGTGCGGCGGT
                        Consensus   (1751) AGAAGCTTGAA CCGG GGA T CG GT CTGGACGTCGGGTG GG GGT
                                           1801                                           1850
              R08E5.1 RNAi frag    (629) GGATCCCATTCTTCGTTGCTCGCCGAGCAATACCCAAAAGCACATTTTGT
rips-1  unspliced (from WB)   (1791) GGATTCCATTCTTCGCTGCTCGCCGAGCAATACCCAAAGTCGCATTTTGT
                        Consensus   (1801) GGAT CCATTCTTCG TGCTCGCCGAGCAATACCCAAA  C CATTTTGT
                                           1851                                           1900
              R08E5.1 RNAi frag    (679) CGGGCTAGAAATTGGAGAAGATGCCATTCGGCAGGCGAAACAACGAAAAA
ips-1   unspliced (from WB)   (1841) CGGGCTAGACATTGGAGAAGATGCCATCCGGCAGGCGAAACAACGCAAGA
                        Consensus   (1851) CGGGCTAGA ATTGGAGAAGATGCCAT CGGCAGGCGAAACAACG AA A
                                           1901                                           1950
              R08E5.1 RNAi frag    (729) CTAAATCAGGAGCGGCTTTCAACAATCTCGAATTTATACAATGCGACGCT
rips-1  unspliced (from WB)   (1891) CTAAATCAGGAGCGGCTTTCAACAATCTCGAATTTATTGAATGTGACGCC
                        Consensus   (1901) CTAAATCAGGAGCGGCTTTCAACAATCTCGAATTTAT  AATG GACGC
                                           1951                                           2000
              R08E5.1 RNAi frag    (779) GGAAAAATGCCAGAAATTTGGACGGATTCCTTCGATTTGGTCCTTATTTT
rips-1  unspliced (from WB)   (1941) GGAAAAATGCCAGAAATCTGGACGGACTCCTTCGATTTGGTGCTTATTTT
                        Consensus   (1951) GGAAAAATGCCAGAAAT TGGACGGA TCCTTCGATTTGGT CTTATTTT
```

**Additional Figure 2B**

*rips-1* mutant TP193 confirmation of causative gene for DTT resistance phenotype.

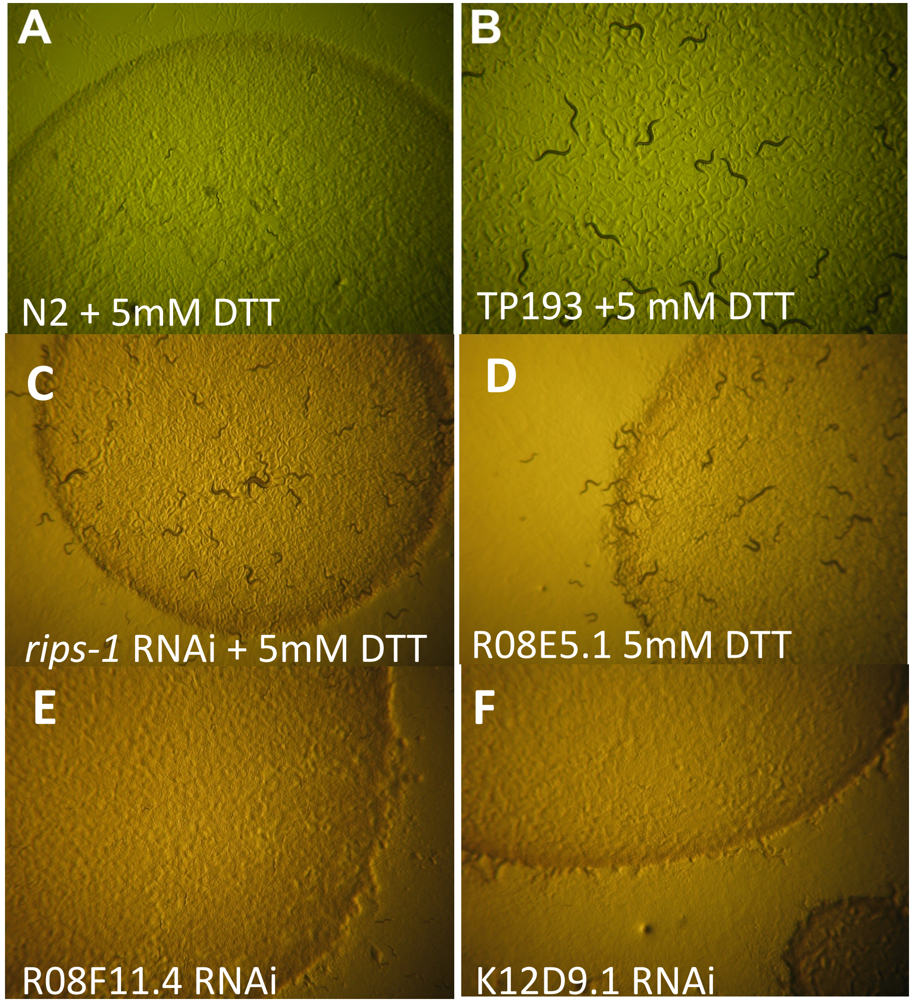


A, N2 arrested development on 5 mM DTT. B, TP193 survival and reaching adult stage on 5 mM DTT. Confirmation of causative gene for DTT resistance phenotype following *rips-1* and homologue RNAi on N2 strain. C, *rips-1* RNAi and survival on 5 mM DTT. D, R08E5.1 RNAi and survival on 5 mM DTT. E, R08F11.4 RNAi and arrest on 5 mM DTT. F, K12D9.1 RNAi and arrest on 5 mM DTT.

**Additional Figure 3**

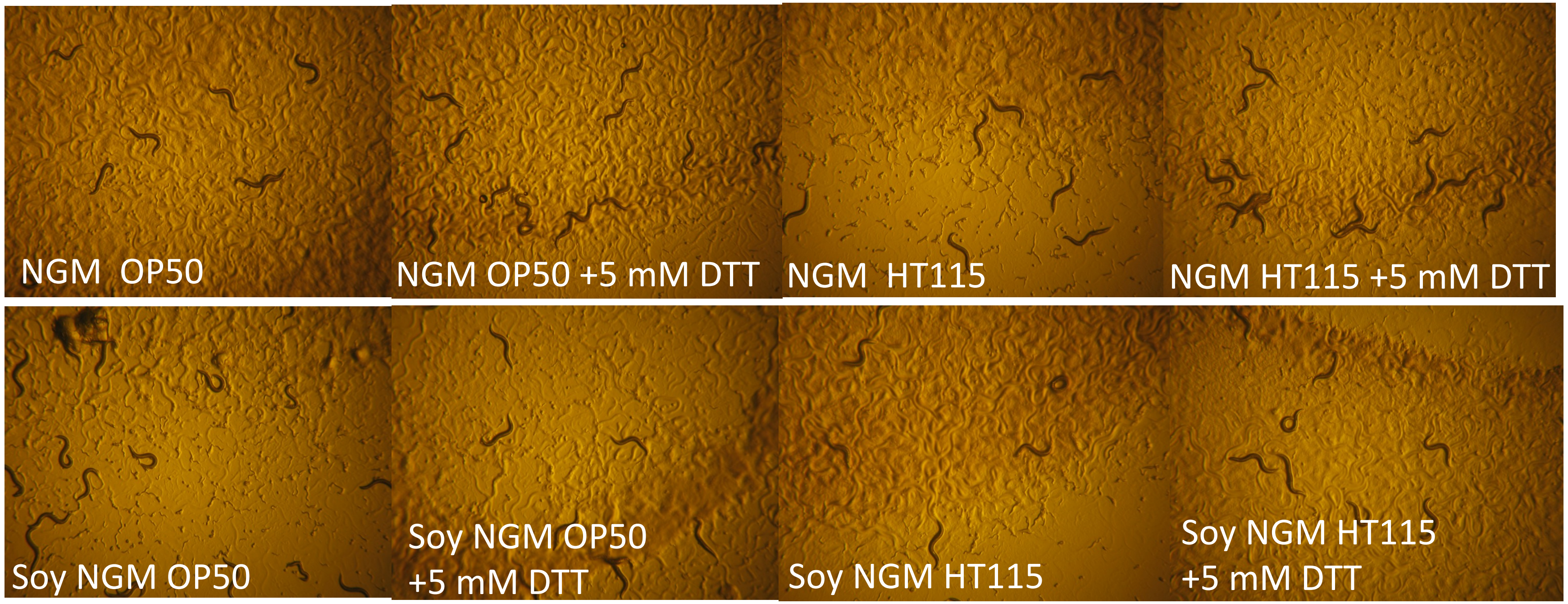


Media composition and bacterial food source do not influence DTT resistance of *rips-1* mutant strain (TP193). TP193 embryos added then viewed as adults after 4 days at 22°C. All growth conditions (animal-based peptone NGM versus Soy plant-based peptone NGM) with different *E. coli* (B12 poor OP50 versus B12 rich HT115) resulted in development to healthy adult populations.

### Additional Figure 4A

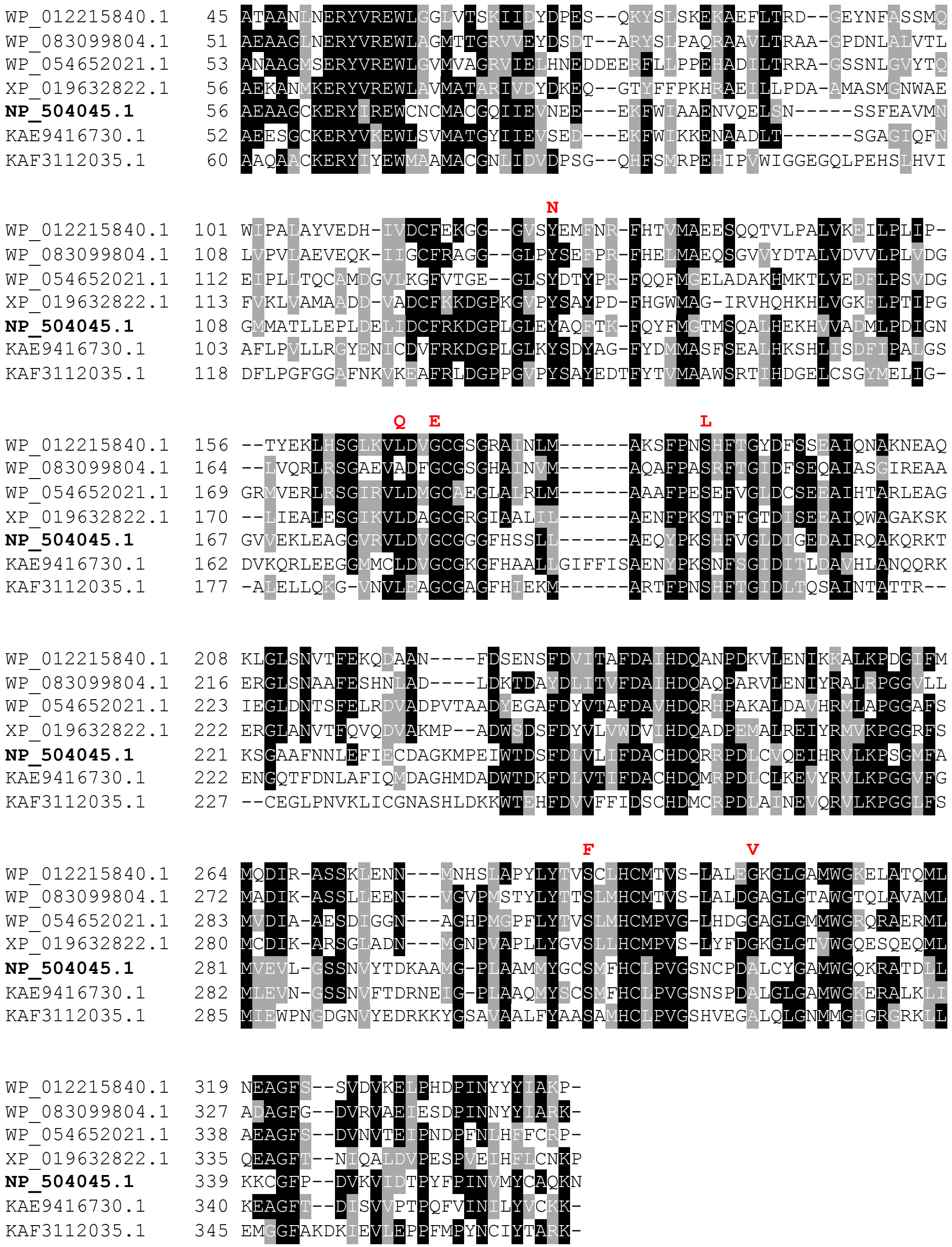


Top BLAST hits from additional File 1 as representative sequences: with conserved mutations found in *rips-1* DDT resistance mutants shown in red above alignments

Archaea [*Nitrosopumilus maritimus*] **WP_012215840.1**

Mycobacterium [*Mycobacterium mantenii*] **WP_083099804.1**

Bacterium [*Desulfovibrio brasiliensis*] **WP_054652021.1**

Non-nematode multi-cellular eukaryote [*Branchiostoma belcheri*] **XP_019632822.1**

*Caenorhabditis elegans rips-1* **NP_504045.1**

Non-Caenorhabditis nematode [*Angiostrongylus cantonensis*] **KAE9416730.1**

Fungi [*Arthrobotrys oligospora*] **KAF3112035.1**

### Additional Figure 4B

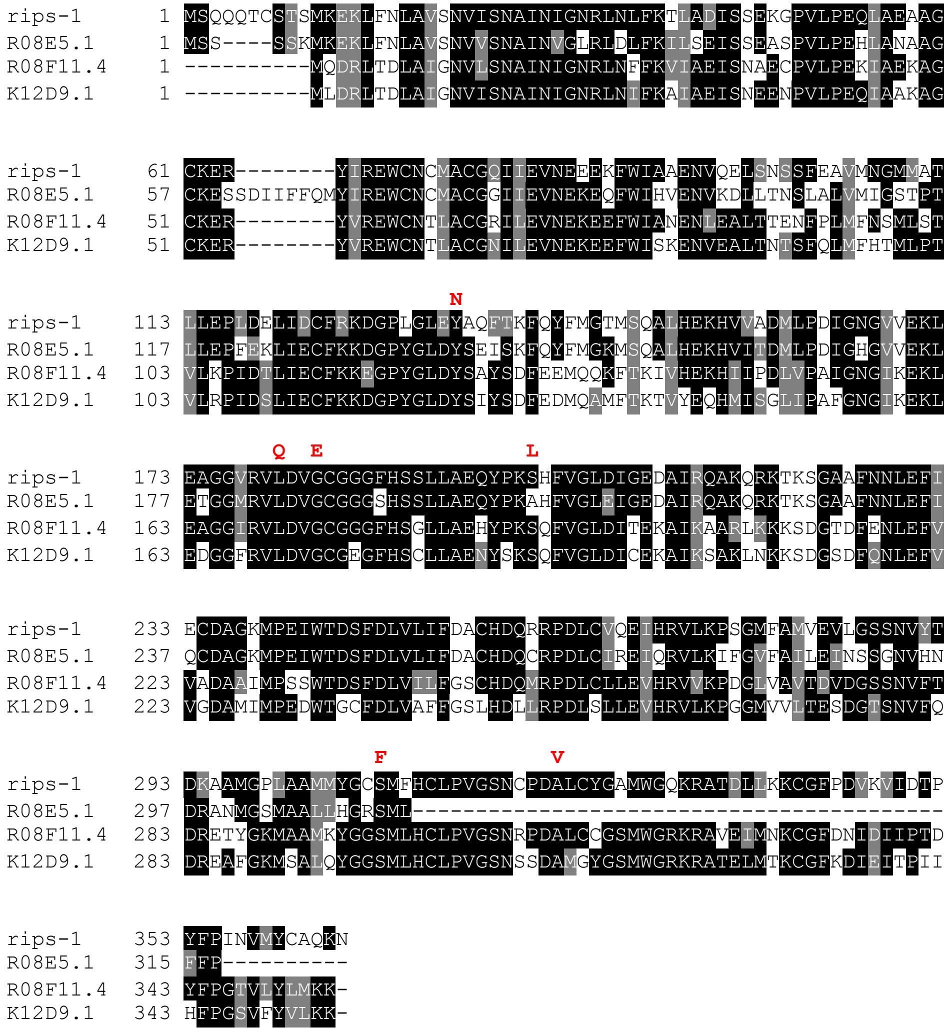


Conservation of amino acids between *rips-1* and orthologues with conserved mutations found in *rips-1* DDT resistance mutants shown in red above alignments. *rips-1* [NP_504045.1], R08E5.1[NP_504044.3], R08F11.4 [NP_504052.1] and K12D9.1 [NP_503823.2].

### Additional Figure 4C

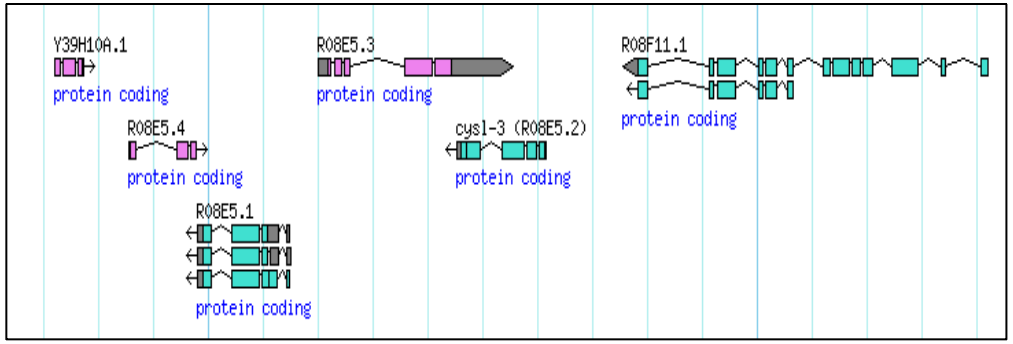


Genomic location and genes surrounding *rips-1* (R08E5.3).

**Additional Figure 5**

RIPS-1::GFP transgenic strain TP313

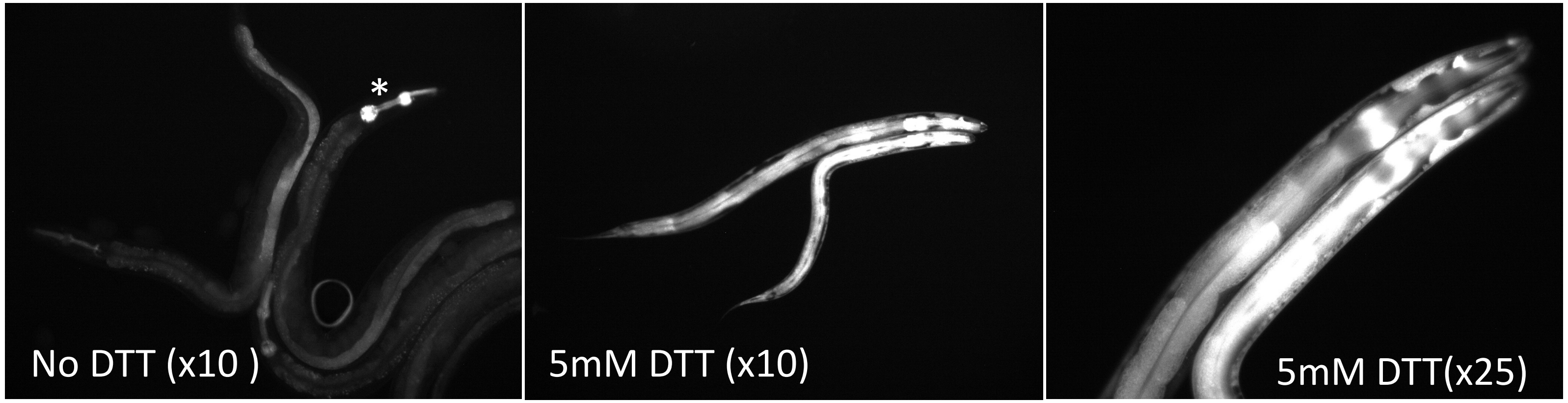


RIPS-1::GFP transgenic strain TP315

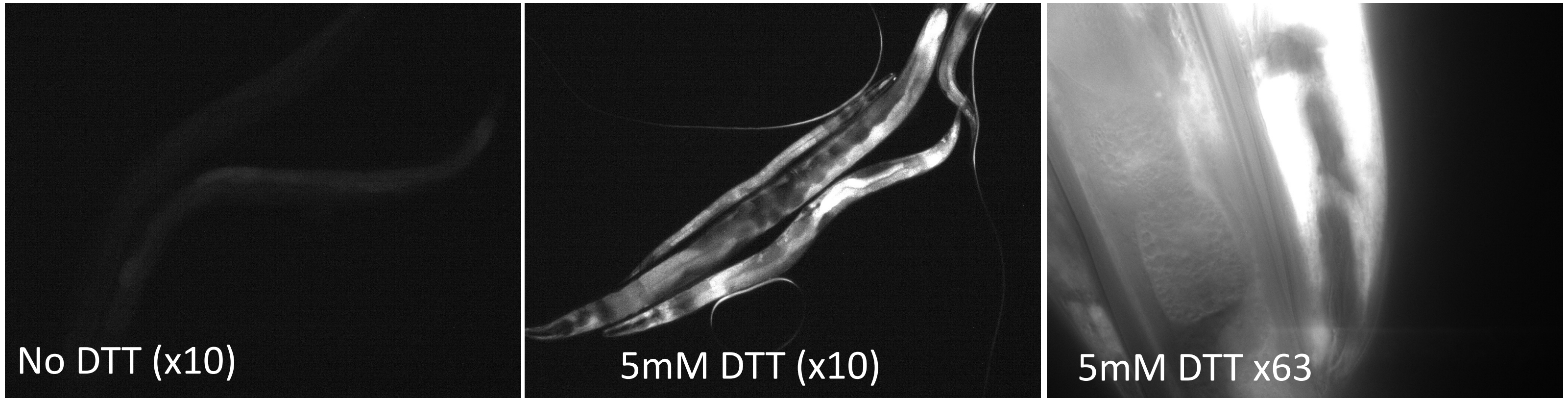


RIPS-1 GFP reporter transgene subcellular induction following 5 mM DTT exposure.
Top panel, transgenic strain TP313, showing weak gut induction in the absence of DTT, (transgenic pharyngeal marker denoted*). Following 5 mM DTT exposure get strong induction in gut and the hypodermis.
Bottom panel, transgenic strain TP315, again showing weak gut induction in the absence of DTT. Following 5 mM DTT exposure get strong induction in gut and the hypodermis.
(Magnification, x10, x25 and x63)

**Additional Figure 6**

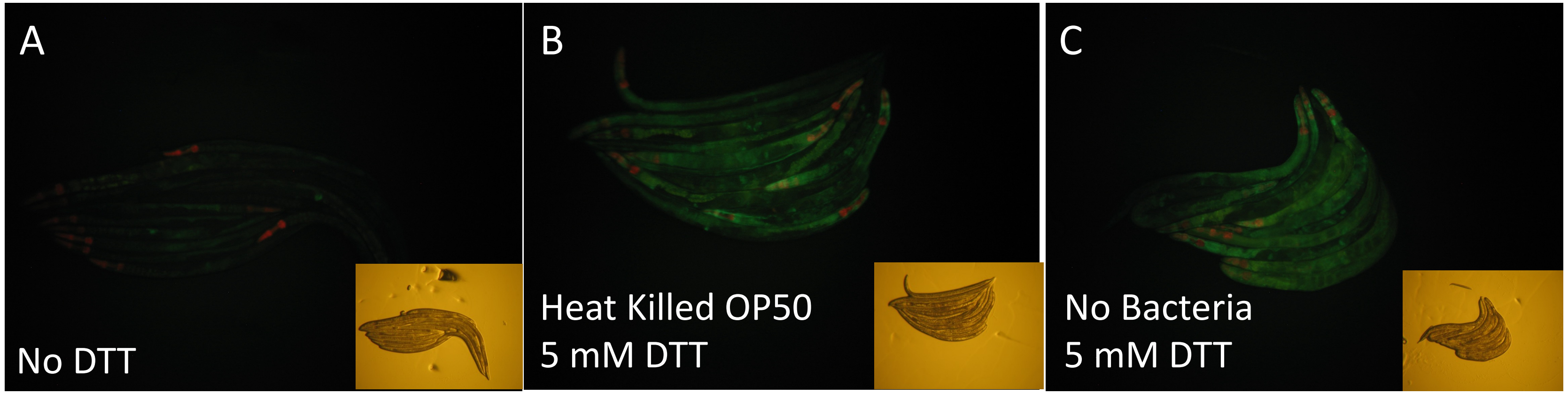


RIPS-1 GFP reporter (TP313) is induced by DTT in the presence of heat killed OP50 (B) and is also induced in the complete absence of bacteria (C). Compares to low induction in absence of DTT (A). Transgenic TP313 nematodes were picked to unseeded plates for 1 hour then transferred to plates in presence or absence of 5 mM DTT, with no bacteria or heat killed bacteria for 24 hours. Bacteria Heat killing involved 60°C for 30 minutes. Insets represent bright field image.

**Additional Figure 7**

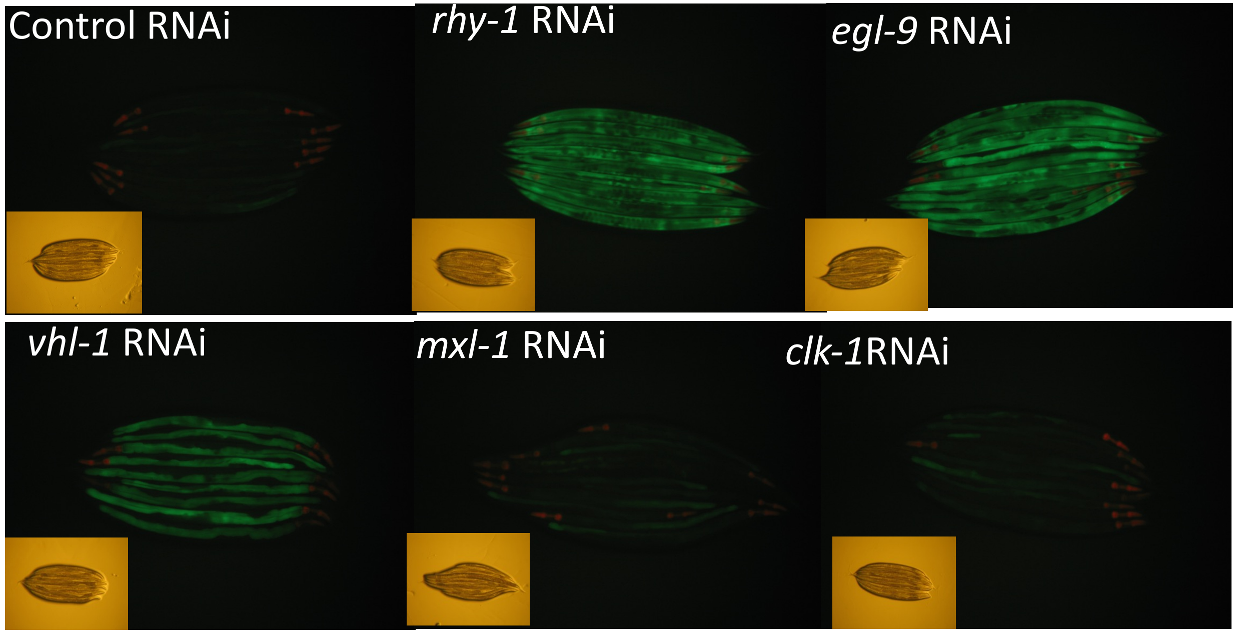


Induction of RIPS-1::GFP reporter following RNAi of TP315. Same experiment depicted in Figure 10 but carried out in independent RIPS-1::GFP reporter strain TP315. Insets represent bright field image.

**Additional Figure 8.**

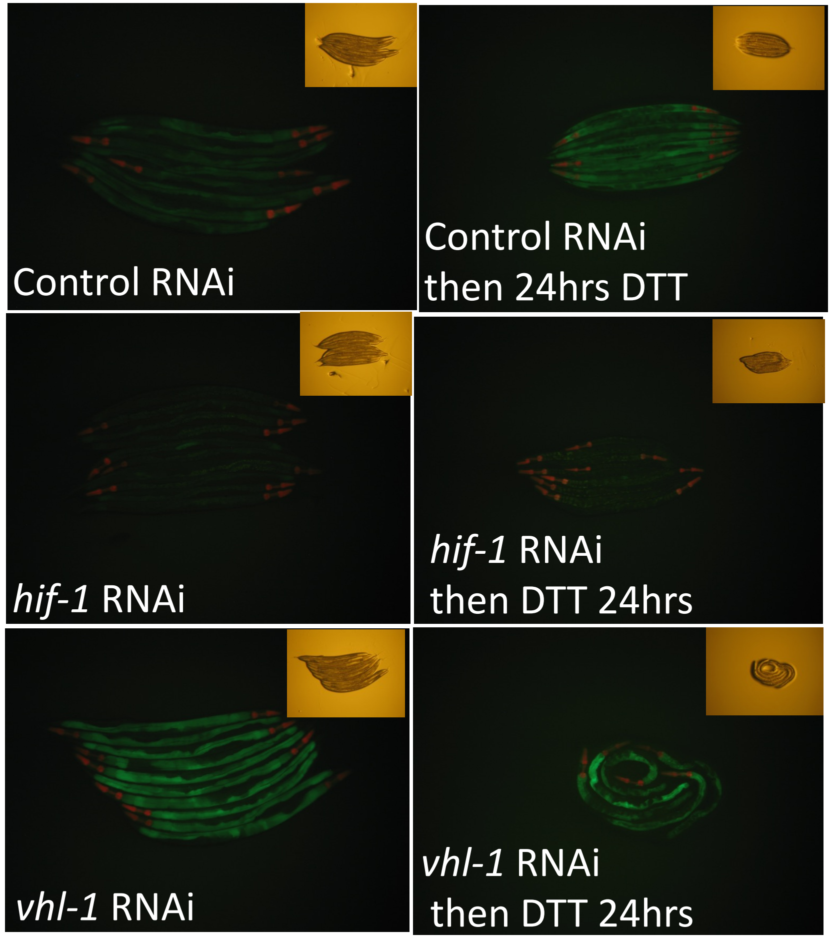


Hypoxia induction factor *hif-1* controls RIPS-1 activation on DTT exposure. Control (empty vector L4440) RNAi, *hif-1* and *vhl-1* RNAi feeding carried out on RIPS-1::GFP reporter (TP315) for 3 days. Then picked *myo-2* transgenic marker (red pharynx) positive L4s to corresponding RNAi plates that were supplemented with 5mM DTT for 24hrs then imaged. Insets represent bright field image.
